# CellProphet: Dissecting Virtual Cell Differentiation through AI-Powered Dynamic Gene Regulatory Network Inference

**DOI:** 10.1101/2025.02.05.636766

**Authors:** Rui Peng, Yuanxu Gao, Yihan Chen, Jinzhuo Wang

**Affiliations:** Department of Big Data and Biomedical AI, College of Future Technology, Peking University, Beijing, 100871, China; Center for BioMed-X Research, Academy for Advanced Interdisciplinary Studies, Peking University, 100871, China; School of Life Sciences, Peking University, Beijing, 100871, China

## Abstract

The AI Virtual Cell (AIVC) framework promises to revolutionize biological research through high-fidelity simulations of cellular behaviors and responses to perturbations. Central to realizing this vision is the ability to model cell differentiation dynamics, which requires accurate inference of gene regulatory network (GRN) that govern cell fate decisions. However, existing computational approaches rely on static GRN models that fail to capture the dynamic changes of regulatory relationships during differentiation, limiting their utility for simulating developmental processes and predicting perturbation outcomes. Here, we present CellProphet, an interpretable AI model that infers dynamic GRN by integrating temporal causality with transformer self-attention mechanism. CellProphet captures time-lagged dependencies between transcription factor (TF) expression and target gene activation while providing interpretable regulatory weights, enabling both accurate prediction and mechanistic insight. When benchmarked against nine state-of-the-art methods across seven differentiation datasets, CellProphet achieves superior performance in all evaluation metrics. Applied to mouse embryonic stem cell differentiation, CellProphet identifies both well-known and potentially novel TFs with substantially high sensitivity and successfully reconstructs dynamic regulatory relations validated through multi-modal epigenomic data. In mouse hematopoietic differentiation, CellProphet accurately predicts cell fate transitions and gene expression changes following *in silico* perturbation of key TFs *Gata1* and *Spi1*, demonstrating its capability for virtual experimentation. These results establish CellProphet as a foundational tool for the AIVC framework, enabling researchers to decode the dynamic regulatory logic of differentiation, accelerate discovery of key regulatory factors, and design targeted cellular interventions for widespread applications.

## Introduction

AI Virtual Cell (AIVC) represents a revolutionary paradigm in computational biology, offering unprecedented capabilities to create comprehensive digital twins of cells that can simulate cellular behaviors and predict responses to perturbations^1^. By learning directly from the exponentially growing omics datasets, AIVC can accelerate discoveries, guide experimental studies, and systematically explore the vast combinatorial space of genetic and pharmacological interventions that determine cellular fate and function. A critical capability of the AIVC framework is simulating the dynamic processes of cell differentiation^1^, the process through which multipotent stem cells transform into the diverse array of specialized cell types. Simulating the differentiation process enables researchers to identify key regulatory factors controlling cell fate decisions, perform *in silico* perturbations on stem cells to predict their cascading effects through developmental pathways, and ultimately engineer desired cell types for regenerative medicine.

The key element of cell differentiation lies the gene regulatory network (GRN), a complex network of interactions between transcription factors (TFs) and their target genes (TGs) that interprets signals, maintains cellular identity, and orchestrates the precise temporal programs guiding cells from pluripotent states toward specialized fates^2^. The GRN serves as the command center that coordinates cellular responses throughout differentiation. Understanding and accurately modeling these regulatory networks is therefore fundamental to simulating differentiation processes within the AIVC framework, as they determine how cells will respond to genetic or pharmacological perturbations.

However, current computational approaches to GRN inference suffer from a critical limitation that fundamentally contradicts the dynamic nature of differentiation. The prevailing methodology involves first clustering cells into discrete types based on their transcriptional profiles, then inferring a separate, static GRN for each identified cluster^3–12^ (**Figure 1A**). These approaches assume that all cells within a type share an identical, fixed regulatory network. This static network paradigm fails to account for a fundamental paradox in developmental biology that the heterogeneous fate outcomes arising from apparently homogeneous cell populations. Specifically, if cells of a given type are governed by identical regulatory architectures, the mechanistic basis for their divergent differentiation trajectories, whereby subpopulations adopt distinct lineage identities under seemingly uniform conditions, remains unexplained. We argue that the GRN itself changes dynamically as differentiation advances, even within the same cell type. Therefore, we require an approach to infer dynamic GRN that can simulate virtual cell differentiation rather than static ones.

**Figure 1.**
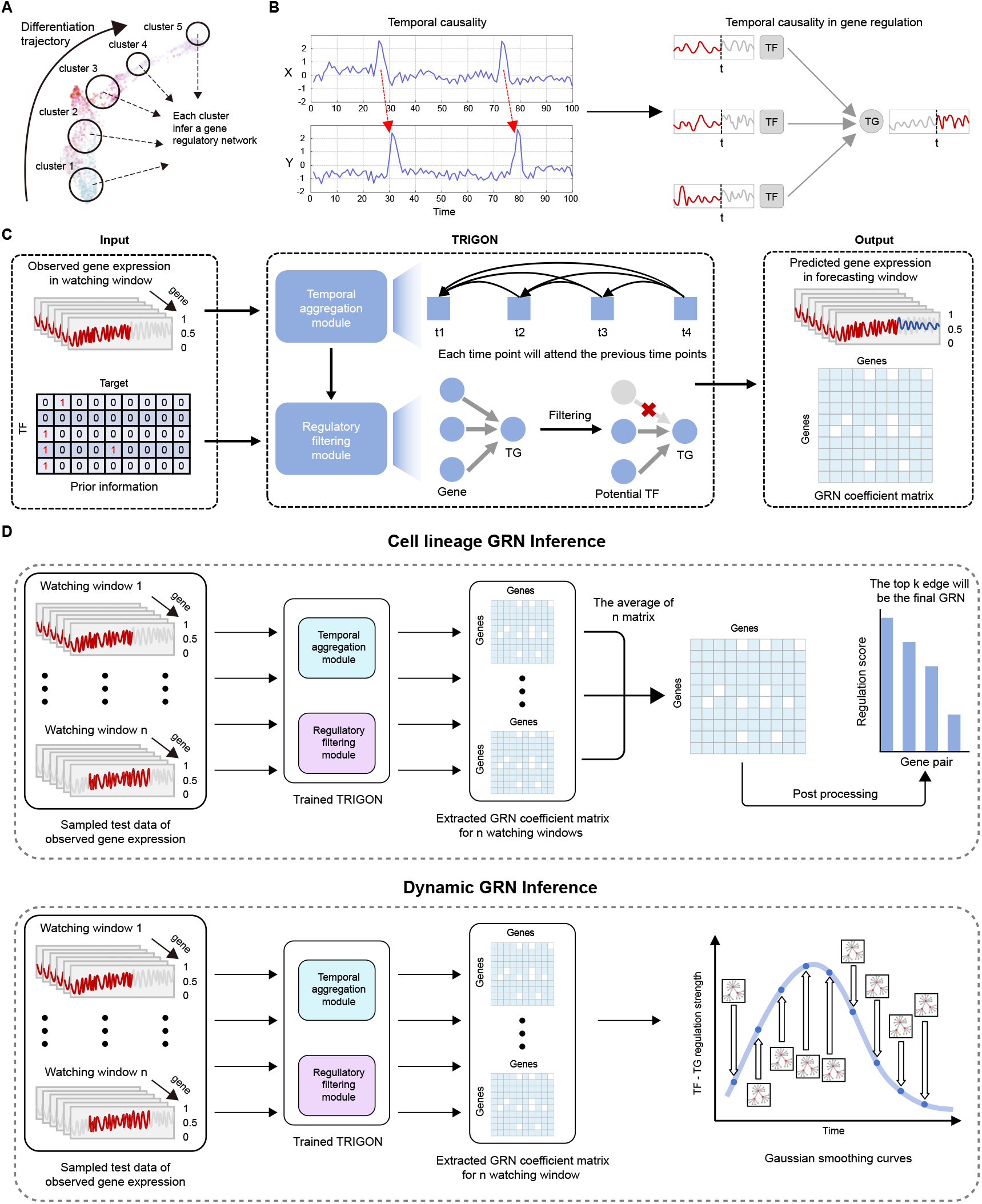
Overview of CellProphet. **A**, Tranditional methods group cells along lineage and construct GRN for each individual cluster, which fail to capture the dynamics of GRN during cellular differentiation. **B**, The concept of temporal causality is observed in gene regulation, where the expression of TG is determined by the expression of its TFs over a given time period. CellProphet leverages this concept to infer dynamic GRN. **C**, CellProphet takes the expression of TFs within the watching window and prior information as input. Through the Temporal Aggregation module and the Regulatory Filtering module, it predicts the expression of TGs in the forecasting window, learns the temporal causality between TFs and TGs and ultimately outputs a GRN coefficient matrix. **D**, The inference procedure for CellProphet begins by sampling test data using the watching window, which yields a sequence of period-specific GRNs. The subsequent processing of these networks is determined by the analytical objective. To infer a single cell lineage GRN, these period-specific GRNs are averaged and then subjected to a final post-processing step. Alternatively, to reconstruct dynamic GRN, a Gaussian smoothing operation is applied across all generated networks.

Beyond capturing dynamics, model interpretability represents another cornerstone requirement for AIVC to achieve its transformative potential. While black-box deep learning models might achieve high predictive accuracy, AIVC must expand the mechanistic understanding of biology by revealing the underlying causal relationships that govern cellular behavior. This dual requirement is essential for AIVC to serve as a tool that narrows the search space for mechanistic hypotheses and accelerates biological discovery.

To address these challenges, we present CellProphet, an AI-powered framework that leverages the advantages of temporal causality and transformer^13^ self-attention mechanisms to infer dynamic GRN for simulating virtual cell differentiation. CellProphet employs temporal causality to capture the inherent time-lagged dependencies between TFs’ expression and downstream target gene activation, enabling the model to learn how regulatory influences propagate through developmental time. Simultaneously, the self-attention mechanism provides interpretable weights that quantify the regulatory influence between gene pairs, allowing researchers to extract biologically meaningful regulatory relationships and identify critical drivers of cell fate decisions.

When benchmarked against nine state-of-the-art GRN inference methods across seven differentiation datasets, CellProphet achieves superior performance in all evaluation metrics with high statistical significance. In mouse embryonic stem cell (mESC) differentiation, CellProphet accurately identifies well-known TFs while uncovering previously unreported regulatory factors potentially involved in lineage specification. Through integration with multi-modal epigenomic data, we validate the authenticity of CellProphet-inferred regulatory relationships and characterize their dynamic changes throughout differentiation trajectories, successfully reconstructing the dynamic GRN underlying mESC lineage commitment. Finally, through *in silico* perturbation experiments targeting the well-characterized TFs *Gata1* and *Spi1* in mouse hematopoietic stem cell (mHSC) differentiation, CellProphet accurately predicts perturbation-induced cell fate transitions and gene expression alterations, demonstrating its capability to fulfill the core promise of the AIVC framework for virtual experimentation and mechanistic discovery. Collectively, these results establish CellProphet as a powerful computational framework for decoding the dynamic regulatory logic of cell differentiation, providing a critical foundation for realizing the transformative potential of AIVC.

## Results

### Overview of CellProphet

CellProphet is a computational framework that leverages transformer and temporal causality to infer dynamic GRN from single-cell transcriptomic data. By combining these two powerful concepts, CellProphet captures the time-dependent regulatory relationships that govern cell differentiation, enabling accurate simulation of developmental processes within the AIVC framework.

CellProphet begins by applying Slingshot^14^ algorithm to assign each cell a specific pseudotime, representing its position along the developmental trajectory. Cells are then ordered according to their pseudotime, transforming the single-cell RNA sequencing^15^ (scRNA-seq) data into a time-series representation of the differentiation process (**Methods**). This temporal ordering captures the sequential gene expression changes that occur as cells progress through differentiation, providing temporal information for each gene across the developmental continuum. With the time-series data constructed, CellProphet integrates the concept of temporal causality to learn gene-specific temporal patterns and infer dynamic GRN. Temporal causality, formalized through Granger causality^16^, determines whether one time series can predict another (**Figure 1B, left**). We posit that this concept naturally applies to gene regulation during differentiation because the expression of a TG at future time points is causally influenced by the expression of its regulatory TFs at earlier time points (**Figure 1B, right**). This time-lagged dependency between TF and TG expression represents the fundamental logic underlying GRN that TFs must be expressed, translated, and localized before they can regulate their targets. To effectively capture these temporal causal relationships, CellProphet employs the transformer’s self-attention mechanism, which not only learns complex temporal dependencies but also provides interpretable attention weights that directly quantify the regulatory strength between gene pairs, offering biological insights into which TFs drive specific expression changes.

CellProphet’s input consists of two components: observed gene expression data and prior information. For the observed gene expression, after assigning pseudotime and ordering cells temporally, we employ two windows to sample the data. For any time point *t*, we sample cells within a watching window of length *w*_1_, spanning [*t,t* + *w*_1_], which contains the observed expression values that serve as model input. We then sample an adjacent forecasting window of length *w*_2_, spanning [*t* + *w*_1_,*t* + *w*_1_ + *w*_2_], which contains the target expression values that CellProphet aims to predict. The model’s core task is to take the observed expression from the watching window and predict the subsequent gene expression in the forecasting window, thereby learning the causal regulatory relationships that drive expression changes over developmental time (**Methods**). The prior information is derived from NicheNet^17^, a comprehensive database that integrates over 50 public data sources for gene regulation information, providing a foundation of established regulatory relationships to guide the inference process.

There are two main modules in CellProphet (**Figure 1C, Extended Figure 1**). The Temporal Aggregation module processes the watching window data by learning historical expression information for each gene. When predicting future gene expression, the module assigns different importance weights to each historical time point within the watching window, recognizing that regulatory influences accumulate over time rather than arising from a single time point (**Methods**). This weighted aggregation captures how multiple past regulatory events collectively shape future expression states, aligning with the fact that gene regulation involves sustained and cumulative effects. The Regulatory Filtering module incorporates the prior information from NicheNet to weight known TF-TG interactions while assigning small, non-zero weights to uncharacterized gene pairs (**Methods**). This ensures that CellProphet leverages existing biological knowledge to improve inference accuracy.

CellProphet produces two outputs. The predicted expression represents the model’s prediction for the forecasting window, while the GRN coefficient matrix quantifies the regulatory strength between each TF-TG pair. The model achieves optimal GRN inference when its predicted gene expression most closely matches the actual observed expression in the forecasting window, as measured by minimizing the mean squared error during training. This design ensures that the inferred GRN accurately captures the regulatory logic driving observed expression dynamics. After training, CellProphet can generate both static lineage-specific GRN (**Figure 1D, up**) through temporal averaging and, critically for AIVC applications, dynamic GRN (**Figure 1D, bottom**) that reveal how regulatory relationships evolve throughout the differentiation process, providing the foundation for high-fidelity virtual cell simulations.

### CellProphet outperforms nine existing methods in accurately inferring GRN across seven benchmarks

To establish CellProphet’s capability for simulating virtual cell differentiation within the AIVC framework, we first conducted comprehensive benchmarking experiments across seven well-characterized differentiation datasets from the BEELINE^18^ benchmark suite. Benchmarking represents a critical validation step for AIVC models, as accurate GRN inference forms the foundation for reliable *in silico* experimentation.

By evaluating CellProphet against established ground-truth networks derived from ChIP-seq data, we assessed whether our temporal causality framework genuinely captures the regulatory logic underlying differentiation.

Testing data for each lineage was sampled based on watching windows. Each window generated a GRN through the inference process, and GRNs from different windows were averaged to construct the final cell lineage GRN (**Figure 2A, Methods**). The accuracy of the inferred GRN can be evaluated using metrics employed in binary classification problems (**Figure 2B, Methods**). We used the area under the receiver operating characteristic curve (AUROC), the area under the precision-recall curve (AUPRC), F1 score, and EPR^18^ as metrics to compare CellProphet against nine latest GRN inference methods: (1) GENIE3^9^ and GRNBoost2^8^, which were widely used classical methods for GRN inference. (2) GRANGER^6^, which was based on Granger causality to infer GRN, (3) CEFCON^10^, NetREX^12^ and Inferelator^7^, which required prior network as input to infer GRN. (4) CellOracle^11^, a multimodal machine learning method leveraged chromatin accessibility data to provide prior knowledge for GRN inference. (5) Random, which randomly selected *k* edges in all potential gene edges (*k*equals the number of edges in the ground truth GRN), and (6) Prior Random, which randomly selected *k* edges in the prior information.

**Figure 2.**
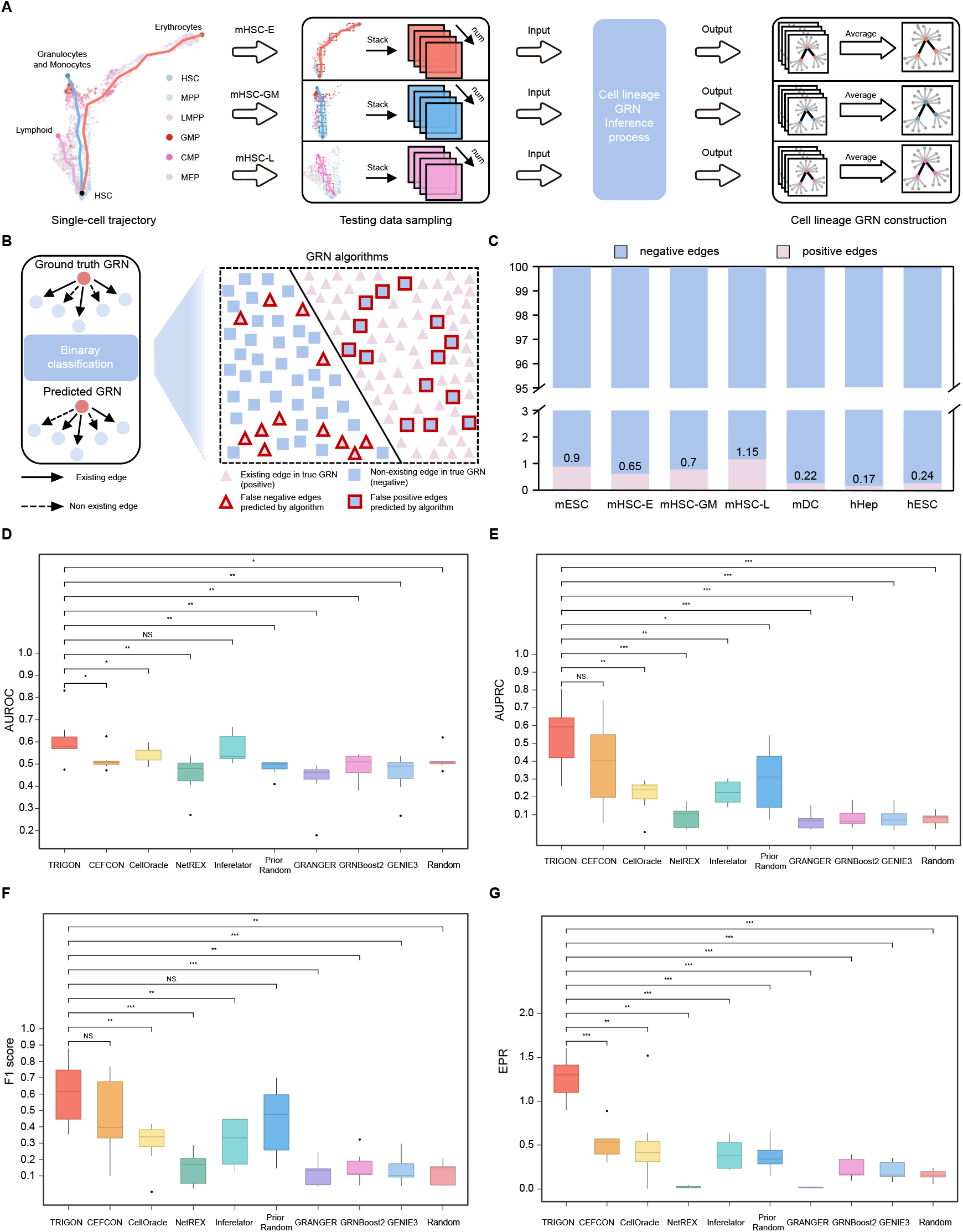
CellProphet shows superior performance over nine methods in seven benchmarks. **A**, CellProphet analyzes each lineage by sampling testing data based on watching windows. GRNs inferred from all windows are averaged, and the top *k* edges are selected to construct the final cell lineage GRN. **B**, GRN evaluation is treated as a binary classification task, determining whether the edges identified by the algorithm are present in the ground truth GRN. **C**, The ratio of positive and negative labels in the seven datasets shows significant class imbalance, with only approximately 1% of edges classified as positive (edges present in the ground truth GRN). **D-G**, Box plots of performance comparisons across seven datasets for AUROC **(D)**, AUPRC **(E)**, F1 score **(F)**, and EPR **(G)**, with statistical significance assessed by Mann-Whitney U test. \**P*<0.05, \*\**P*<0.01, \*\*\**P*<0.001, NS. Not significant.

The average AUROC of all benchmark datasets (**Figure 2D, Extended Table 2**) indicated that CellProphet achieved the highest performance with an AUROC of 0.61, significantly surpassing most methods. We observed that previously proposed methods showed no significant performance improvement compared to Random on this metric. For instance, on the mESC dataset, CellOracle (0.597), GRNBoost2 (0.537), and GENIE3 (0.545) performed only marginally better than Random (0.505), indicating that the accuracy of these methods in GRN inference is barely above random guessing. However, CellProphet achieved an AUROC of 0.831, demonstrating the model’s superior performance. Due to a pronounced class imbalance in label distribution, with positive labels accounting for less than 2% across all datasets (**Figure 2C, Extended Table 1, Methods**),we adopted AUPRC for performance evaluation, as it was a superior metric compared to AUROC in this scenario^19^. In that evaluation **(Figure 2E, Extended Table 2)**, CellProphet achieved an average AUPRC of 0.541 across seven datasets, representing a 162% improvement over the latest method, CellOracle (0.206). To further validate CellProphet’s performance, we additionally calculated the F1 score and EPR across all datasets. Statistical tests across the seven benchmark datasets confirmed that CellProphet’s F1 score and EPR values were also significantly higher than those of nearly all competing methods **(Figure 2F-G, Extended Table 2)**. To determine whether prior knowledge accounted for CellProphet’s advantage, we carried out a supplementary evaluation in which the prior information was removed. Algorithms that cannot run without a prior network (CEFCON, Celloracle, NetREX, Inferelator, and Prior Random) were therefore excluded, leaving CellProphet, GRANGER, GRNBoost2, GENIE3, and Random for a strictly de novo comparison. Even in this setting, CellProphet demonstrated state-of-the-art performance across all evaluation metrics on all datasets, underscoring its robustness to the absence of prior information (**Extended Figure 3, Extended Table 3**).

### CellProphet identifies well-established and potentially novel TFs governing cell fate decisions

Identifying key TFs that orchestrate cell fate decisions represents a fundamental capability for virtual cell simulation. Data-driven identification of regulatory factors enables researchers to predict which genetic perturbations will redirect differentiation trajectories, design targeted interventions for cellular reprogramming, and discover novel therapeutic targets for regenerative medicine. To evaluate the model’s ability to identify relevant TFs, we applied CellProphet to the mESC dataset (**Figure 3A**), which comprised scRNA-seq data collected at five distinct time points (0, 12, 24, 48, and 72 hours) as mouse embryonic stem cells progress toward primitive endoderm (PrE). We utilized AUCell^19^ (**Extended Figure 2, Methods**) to calculate the activity score of each TF regulon in individual cell based on the inferred GRN. A TF regulon was defined as a gene set comprising the TF itself and its TGs (**Figure 3B**). The activity score reflected the proportion of genes within the TF regulon that were highly expressed in a given cell, with a higher score indicating that the TF regulon was more active in that cell. We performed Analysis of Variance (ANOVA) to compare the activity score of each TF’s regulon across different time points and selected those with significant differences (p-value < 0.01) as the identified key TFs (**Methods**). In the mESC dataset, 37 key TFs were identified by CellProphet (**Figure 3C**), and the dynamic changes in their activity scores were visualized in **Extended Figure 4**.

**Figure 3.**
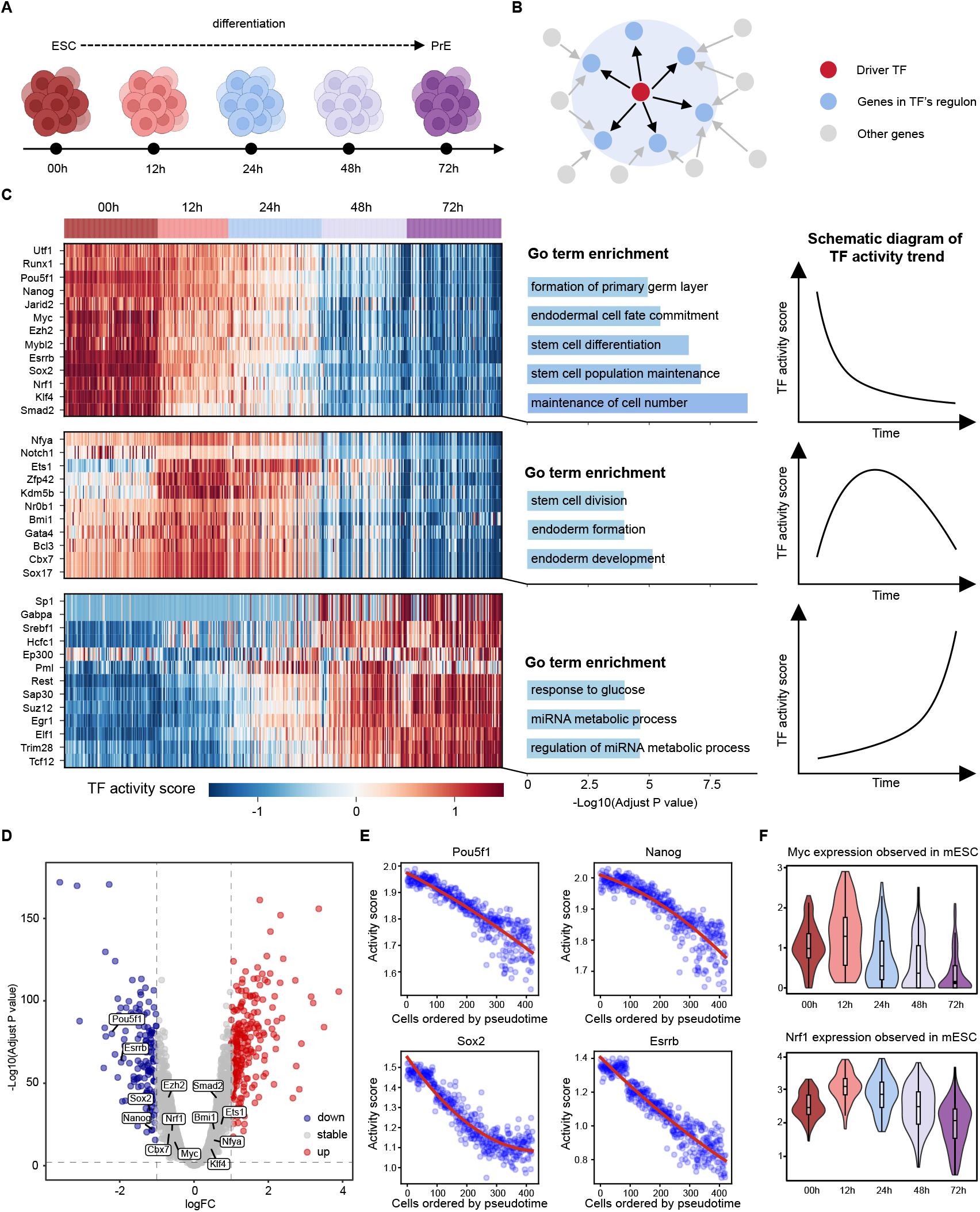
CellProphet is capable of identifying key TFs involved in the process of cellular differentiation. **A**, The mESC dataset consists of 421 scRNA-seq data capturing the differentiation of mouse embryonic stem cells into primitive endoderm, sequenced at five distinct time points. **B**, A TF regulon is defined as a gene set comprising the TF itself and its target genes, as determined by the inferred GRN. **C**, The activity score of each TF regulon is calculated using the AUCell algorithm. ANOVA is performed on TF activity scores across the five sequencing time points, and TFs with p-value < 0.01 are identified as key TFs. The line plots at right are schematic diagrams that illustrate the TF-activity trends observed in the heatmaps. **D**, Volcano plot from DEA, where the logFC threshold for identifying key TFs is set to 1, and p-values are corrected using Bonferroni correction. DEA identifies a subset of key TFs (e.g., *Pou5f1, Nanog* and *Sox2*) also detected by CellProphet, while a group of genes labeled as “stable” could not be identified as key TFs. **E**, The activity scores for key TFs identified by CellProphet are fitted with quadratic curves, illustrating their dynamics over differentiation process. The x-axis represents the cells ordered by pseuditime. As each cell is assigned a temporal value, this axis can be interpreted as the progression of time. **F**, Violin plots show the original expression of two key TFs identified by CellProphet but not by DEA. These TFs exhibit minimal changes in expression across the five sequencing time points.

CellProphet identified three categories of key TFs based on the temporal dynamics of their activity scores during differentiation. (1) Early-stage TFs: These TFs exhibited high activity scores at the onset of differentiation (00h) and gradually declined as differentiation advanced. These TFs were highly expressed during the early stages of differentiation, suggesting their involvement in maintaining stem cell identity. This was further supported by GO term enrichment analysis, which linked them to “stem cell number maintenance” and “stem cell population maintenance”. Among them, *Nanog, Pou5f1, Esrrb*, and *Sox2* were well-established regulators of stem cell pluripotency^20,21^, *Utf1* and *Ezh2* have been previously reported to play critical roles in maintaining stem cell identity^22–24^. However, TFs like *Nrf1, Klf4, Runx1*, etc., these TFs have not yet been reported to be associated with early mESC development. They are the potential novel TFs identified by CellProphet warranting further experimental validation. (2) Mid-stage TFs: This group was characterized by transient increases in activity scores during the intermediate stages of differentiation (12h and 24h), suggesting their roles in initiating and promoting the differentiation process. GO term enrichment of these genes highlighted terms related to “embryonic development”. Among them, *Gata4* was reported to direct the development of two distinct types of Sox17+ endoderm, including an epCam+Dpp4+ subtype of visceral endoderm^25^. (3) Late-stage TFs: These TFs exhibited high activity scores during the terminal stages of differentiation (48h and 72h). Intriguingly, GO enrichment analysis revealed these TFs were significantly associated with “metabolic process” term. For instance, *Srebf1*, a well-known master regulator of fatty acid metabolism^26^, has not been previously implicated in mESC differentiation. These metabolic-related TFs were potential TFs identified by CellProphet, they exhibited marked upregulation in their activity score at the terminal stage of differentiation. This finding suggested a metabolic shift may represent an integral yet underappreciated component of stem cell differentiation, highlighting a promising direction for future research.

To verify CellProphet’s advantage over differential expression analysis (DEA) in identifying TFs, we calculated the log fold change (logFC) in gene expression and identified the key TFs based on p-value (**Figure 3D**). DEA highlighted four TFs (*Pou5f1, Nanog, Sox2*, and *Esrrb*), whose expression decreased during differentiation. This trend was consistent with the TF activity scores calculated by CellProphet (**Figure 3E**). However, CellProphet identified a group of genes (*Klf4, Myc, Nrf1*, etc.) that could not be distinguished by DEA. They were labeled as “stable” in DEA and were not identified as key TFs. Violin plots of *Myc* and *Nrf1* expression across different time points further supported this, as their expression showed minimal changes that can’t be detected by DEA (**Figure 3F**). In contrast, these genes were recognized as key TFs by CellProphet, with their activity scores being high during the early stages of differentiation (**Figure 3C**). This finding was consistent with previous reports that *Myc* and *Nrf1* promoted the maintenance of stem cell pluripotency^27,28^, showing the capacity of CellProphet to identify TFs with greater precision.

### CellProphet’s predicted target genes are supported by epigenomic data

Having established CellProphet’s capability to identify key TFs governing differentiation, we next sought to validate the authenticity of specific TF-TG regulatory relationships inferred by our model. Accurate prediction of regulatory edges represents a more stringent test of GRN inference methods, as it requires not only identifying important regulators but also correctly mapping their downstream targets. This validation is essential for AIVC applications, where simulating perturbation effects depends on accurately modeling how TFs influence their target genes. Without validated TF-TG relationships, virtual cells cannot reliably predict cascading effects of genetic interventions.

To rigorously assess the biological validity of CellProphet-inferred regulatory edges, we focused on *Nanog*, a well-known TF of pluripotency during mESC differentiation. From CellProphet’s inferred GRN, we extracted *Nanog*’s sub-regulatory network and identified its top-scoring predicted targets (**Figure 4B**). We selected four high-confidence targets *Eps8, Fkbp4, C2cd5*, and *Ptpn6* for detailed validation. These genes were chosen based on three criteria: (1) genomic proximity to *Nanog* (within 60 Mb), enabling Hi-C validation of physical interactions. (2) ranking in the top 20% of regulatory strength scores. and (3) availability of comprehensive multi-omics validation data.

**Figure 4.**
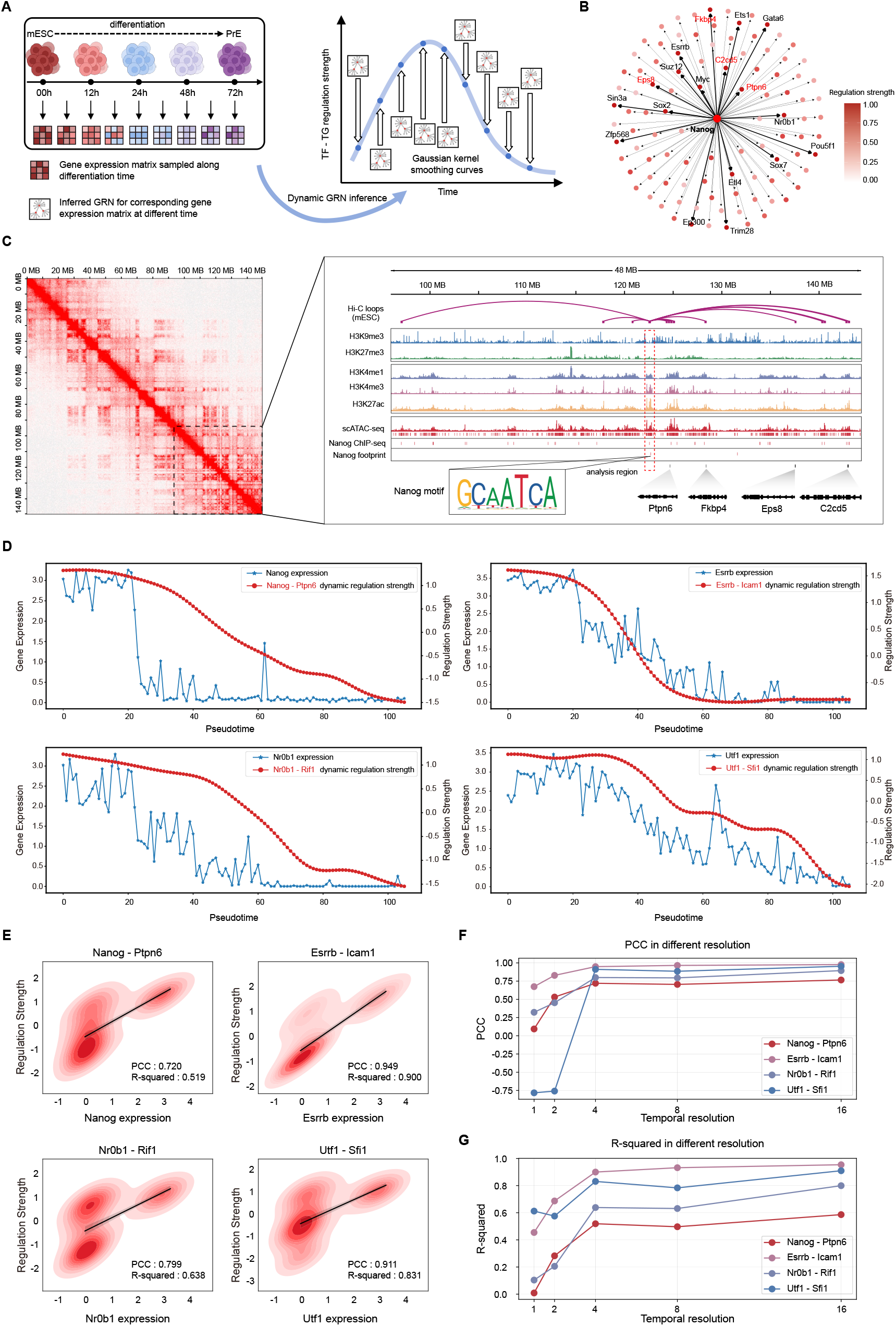
CellProphet identifies target genes and constructs dynamic GRN. **A**, Pipeline of dynamic GRN construction. For each watching window, the TF-TG regulatory strengths inferred in the GRN are smoothed using Gaussian smoothing to generate temporal variation curves. **B**, The regulatory network of *Nanog* inferred by CellProphet, with the regulatory strengths between *Nanog* and its target genes highlighted using color intensity. **C**, Epigenomic data for mouse embryonic stem cells, including histone modification, scATAC-seq, and Hi-C data, provide evidence that CellProphet correctly identifies the regulatory relationships between *Nanog* and its four target genes (*Ptpn6, Fkbp4, Eps8*, and *C2cd5*). **D**, Dynamic GRN inferred at temporal resolution of 4 demonstrate that the regulatory dynamics of *Nanog, Esrrb, Nr0b1*, and *Utf1* align closely with their gene expression trends. **E**, KDE plots of TF expression versus dynamic regulatory strength show PCC above 0.7, with *Esrrb* and *Utf1* achieving PCC as high as 0.9, indicating a strong correlation between changes in gene expression and TF regulatory dynamics. **F-G**, CellProphet analyzes dynamic network changes across different temporal resolutions. Networks inferred at a resolution of 1 show the lowest performance. As the resolution changes to 16, the PCC and R-squared steadily improve.

To validate these predicted regulatory relationships, we integrated supporting evidence from a previously published study^29^, which provided high-throughput chromosome conformation capture (Hi-C) data^30^, histone modification data (H3K9me3, H3K27me3, H3K4me1, H3K4me3, H3K27ac), single-cell assay for transposase-accessible chromatin sequencing (scATAC-seq) data^31^, and chromatin immunoprecipitation sequencing (ChIP-seq) data. Hi-C data were processed into a chromatin-contact matrix and displayed as a heatmap centered on *Nanog* (**Figure 4C, left, 90-150 Mb**). Within this window, we identified Hi-C loops, ATAC peaks and a *Nanog* footprint **(Methods)**. We then overlaid these features with histone-mark and ChIP-seq tracks^32^ and visualized in Integrative Genomics Viewer^33^ (IGV) **(Figure 4C, right)**. We focused on the region containing *Nanog* binding motifs. Within this region, ATAC-seq data revealed prominent chromatin accessibility peaks, indicating open chromatin states conducive to transcription factor binding. This region showed enrichment for activating histone marks (H3K4me1/3, H3K27ac) and depletion of repressive marks (H3K9me3, H3K27me3), collectively confirming this region as active regulatory elements. Critically, Hi-C loops connected these *Nanog*-bound regulatory elements to the genomic loci of *Eps8, Fkbp4, C2cd5*, and *Ptpn6*, the exact *Nanog* targets predicted by CellProphet. These evidences (*Nanog* binding motifs, chromatin accessibility, activating histone modifications, and three-dimensional chromatin interactions) provided robust multi-modal validation that CellProphet not only identifies key TFs but also accurately maps their downstream regulatory targets, proving its capability to reconstruct genuine biological regulatory networks.

### CellProphet reconstructs dynamic regulatory network for identified TF-TG interaction

After validation of CellProphet’s ability to identify key TFs and their corresponding targets, we further tested our model’s ability to infer dynamic GRN. During differentiation, the regulatory strength between TFs and their TGs changes continuously, some connections strengthen while others weaken or disappear entirely. This dynamic change orchestrates cells differentiate into specific lineages.

To demonstrate CellProphet’s dynamic GRN inference capability, we tracked how regulatory relationships of key factors evolved throughout mESC differentiation. In contrast to our approach for static lineage GRN construction, we applied Gaussian smoothing across inferred GRN coefficient matrices to capture continuous regulatory dynamics (**Figure 4A, Methods**). A key feature of CellProphet is its flexible temporal resolution, which we define as the number of consecutively ordered cells in the watching window. By adjusting this parameter, we can analyze regulatory dynamics at different biological scales. For instance, a watching window of 1 cell provides single-cell resolution for detecting transient regulatory states, while larger windows (e.g., 16 cells) yield population-level trends with greater statistical robustness. This multi-scale capability is essential for investigating diverse biological questions, from rapid cell fate decisions to stable developmental programs.

We selected four well-characterized factors (*Nanog, Esrrb, Nr0b1*, and *Utf1*) along with their validated target genes to track regulatory strength changes throughout mESC differentiation. This analysis was based on the fundamental premise that increased expression of a TF should correlate with enhanced regulatory strength toward its target genes, implying that the dynamic expression profile of a TF should align with the inferred trajectory of its regulatory interactions. We visualized the dynamic changes in TF–TG regulatory strength, as predicted by CellProphet, alongside the actual gene expression dynamics of the corresponding TFs (**Figure 4D**). At a temporal resolution of 4 cells, the regulatory strengths of these TFs exhibited a progressive decline throughout differentiation, a trend consistent with the reduction in their expression levels. To quantitatively assess this concordance, we computed the Pearson correlation coefficient (PCC) between each TF’s expression and its predicted regulatory strength (**Figure 4E**). All correlations exceeded 0.7, with *Esrrb* and *Utf1* reaching PCC as high as 0.9. This strong agreement demonstrates CellProphet’s capability to accurately infer dynamic GRN.

To evaluate the tradeoff between temporal resolution and inference robustness, we analyzed the same regulatory relationships at resolutions ranging from single-cell (1 cell) to population-level (16 cells) (**Figure 4F-G, Extended Figure 5**). Notably, single-cell resolution exhibited the lowest correlation between inferred regulatory strength and TF expression, whereas population-level resolution achieved the highest correlation. This pattern is likely attributable to the inherent sparsity of scRNA-seq data, which stems from limited sequencing depth and results in widespread “dropout” events (i.e., a substantial proportion of zero counts reflecting technical artifacts rather than true biological absence). Consequently, GRN inferred from individual cell is particularly prone to technical artifacts and sampling bias. As the number of cells increased, these technical artifacts were progressively averaged out, leading to more robust and accurate reconstruction of dynamic regulatory network. A similar phenomenon was observed for another well-known TF *Sox2* (**Extended Figure 6**), at temporal resolution of 16 cells, CellProphet achieved a PCC of approximately 0.8.

The ability to reconstruct dynamic GRN at multiple temporal resolutions provides crucial flexibility for different AIVC applications. High-resolution dynamics can reveal transient regulatory states during cell fate decisions, while population-level analysis captures robust developmental programs. This multi-scale capability, combined with validated regulatory relationships, establishes CellProphet as a powerful tool for simulating the dynamic regulatory processes underlying cellular differentiation in virtual cell models.

### CellProphet enables *in silico* perturbation to predict cell fate changes

Following the validation of CellProphet’s capacity to identify key TFs, map their regulatory targets, and capture dynamic regulatory changes throughout differentiation, we proceeded to evaluate its performance in predicting cell responses to genetic perturbations, the central objective of the AIVC framework. Accurate *in silico* perturbation requires not only correct GRN topology but also precise quantification of regulatory strengths, as these determine how perturbation effects propagate through the network to alter cell fate. The ability to accurately simulate cell responses to perturbations offers significant practical value, allowing researchers to computationally predict phenotypic outcomes, identify effective intervention strategies, and design reprogramming protocols, thereby reducing reliance on large-scale experimental screening.

We employed a method introduced by CellOracle^11^ to evaluate whether the predicted GRN could recapitulate the fate changes following the knockout of key TFs. The GRN inferred by CellProphet was represented as a directed graph, with edges indicating regulatory strengths from TFs to TGs. Perturbing a specific TF propagated the effects to its TGs along these edges, where the extent of the effect on each TG was determined by the edge weight (**Figure 5A**). To simulate gene knockout, the expression of the perturbed TF was set to zero, resulting in a perturbed expression vector. This vector was subsequently multiplied by the GRN coefficient matrix, propagating the perturbation effects to the TGs (**Extended Figure 2, Methods**). By iteratively propagating these effects, we can finally computed a perturbation vector for each cell (**Extended Figure 2, Methods**). These vectors indicated the direction of fate change for each cell following the perturbation (**Figure 5A, Methods**).

We applied CellProphet to perform *in silico* perturbation on the mHSC dataset, which has several key cellular states: (1) hematopoietic stem cell (HSC), (2) multipotent progenitor (MPP), (3) lymphoid multipotent progenitor (LMPP), (4) common myeloid progenitor (CMP), (5) megakaryocyte-erythrocyte progenitor (MEP), and (6) granulocyte-monocyte progenitor (GMP) (**Figure 5B**). We focused on two well-characterized TFs, *Gata1* and *Spi1*, to investigate cell fate changes following *in silico* knockout. Previous studies have demonstrated that *Gata1* promoted erythrocytes lineage development in HSC and facilitated the differentiation of CMP into MEP^35–37^. In contrast, *Spi1* enhanced LMPP differentiation and promoted the differentiation of CMP into GMP while simultaneously inhibiting their differentiation into MEP^38^ (**Figure 5C**).

**Figure 5.**
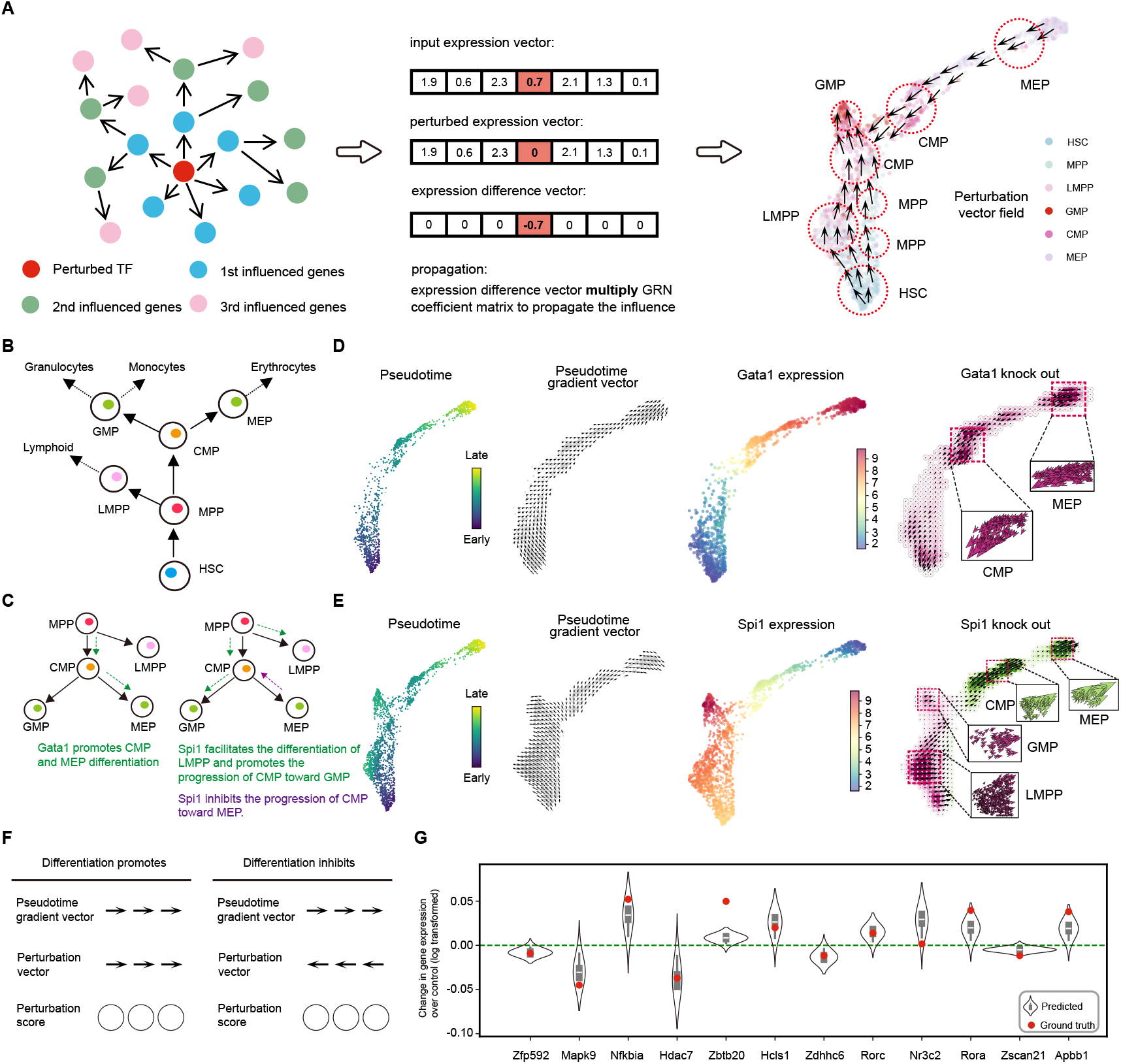
CellProphet predicts cell fate changes after *in silico* perturbation. **A**, Pipeline of *in silico* perturbation. The expression of perturbed TF is set to zero, generating an expression difference vector. This vector is multiplied by the inferred GRN coefficient matrix to propagate the perturbation’s effects to the target genes of the TF. Each cell ultimately generates a perturbation vector, whose direction indicates the cell’s fate changes following the perturbation. **B**, The three lineages of mHSC dataset, detailing the associated cell states and differentiation trajectories. **C**, *Gata1* promotes the differentiation of HSC toward the erythrocytes lineage, while *Spi1* promotes the differentiation of HSC toward granulocytes-monocytes lineage and lymphocytes lineage, and inhibits erythrocytes lineage differentiation. **D**, Pseudotime and *Gata1* expression are colored on the erythrocytes lineage, along with the pseudotime gradient vector field and *Gata1* knockout simulation vector field annotated with perturbation scores. **E**, Pseudotime and *Spi1* expression are colored for the three lineages, along with pseudotime gradient vector fields and *Spi1* knockout simulation vector fields annotated with perturbation scores. **F**, Perturbation score calculation process. If the pseudotime gradient vector and the perturbation vector are aligned in the same direction, the score is positive; if they are opposite, the score is negative. **G**, CellProphet predicted gene expression changes following *Gata1* knockout, with the top 12 predicted genes selected. The ground truth is derived from the experimental difference between perturbed and wild-type cells.

We conducted *in silico* perturbation of *Gata1* in the erythrocytes lineage (**Figure 5D**). Each cell was colored based on its pseudotime, and the pseudotime gradient vector was computed (**Methods), which indicated the natural direction of differentiation**. By visualizing *Gata1* expression, we observed high expression in CMP and MEP, consistent with its role in erythrocytes specification. Upon *in silico* knockout of *Gata1* (**Methods**), we calculated the perturbation vector for each cell and found that these vectors were opposite to the actual differentiation trajectory. Notably, perturbation vectors were more densely clustered in regions containing CMP and MEP, the cells that have high *Gata1* expression. To quantify the effect of *Gata1* perturbation, we calculated the perturbation score for each cell (**Methods**). If the pseudotime gradient vector and perturbation vector had the same direction, the perturbation score was positive, and cells were colored green, indicating that the TF knockout promoted the cell’s differentiation. Conversely, the perturbation score was negative, and cells were colored purple, indicating that the TF knockout inhibited the cells’ differentiation (**Figure 5F**). Upon *Gata1* knockout, all cells in the erythrocytes lineage were colored purple (**Figure 5D**), showing a widespread inhibition of differentiation. The perturbation scores were particularly pronounced in regions containing CMP and MEP, demonstrating a greater impact on these populations, which was consistent with the high expression of *Gata1* in these cell types. Next, we conducted *in silico* knockout of *Spi1* across cells from all three lineages. We visualized pseudotime, *Spi1* expression, and the pseudotime gradient vectors for all cells (**Figure 5E**). *Spi1* was found to be highly expressed in LMPP and GMP, with low expression in MEP. Following *Spi1* knockout, the perturbation vectors for LMPP and GMP were directed toward MEP, and these cells received negative perturbation scores, indicating that *Spi1* knockout suppressed their differentiation and caused them to progress toward MEP. CMP exhibited perturbation vectors directed toward MEP and received positive perturbation scores, indicating that *Spi1* knockout promoted their differentiation into MEP. These findings are consistent with the known biological role of *Spi1*, which promoted the differentiation of GMP and LMPP, but suppressed the differentiation of CMP toward the MEP lineage (**Figure 5C**). Thus, following *Spi1* knockout, MEP differentiation was enhanced, while the development of GMP and LMPP was inhibited. To further validate prediction accuracy, we compared CellProphet’s predicted gene expression changes following *Gata1* knockout with actual experimental data. CellProphet successfully predicted the expression alterations of key downstream genes, demonstrating its ability to forecast not only cell fate transitions but also molecular-level responses to perturbations (**Figure 5G**).

These *in silico* perturbation experiments demonstrate that CellProphet captured the causal regulatory logic governing cell fate decisions with sufficient accuracy to predict perturbation outcomes, achieving the core function of AIVC.

## Discussion

The development of AIVC represents an ambitious vision to create comprehensive digital twins capable of simulating cellular behaviors, predicting responses to perturbations, and accelerating biological discovery. Among the diverse cellular processes that AIVC aims to capture, simulating cell differentiation represents a particularly critical capability. Differentiation underlies development, tissue homeostasis, and regenerative medicine, making its accurate modeling essential for applications ranging from understanding congenital diseases to engineering cell-based therapies. Virtual differentiation experiments could reveal how stem cells navigate toward specialized fates, identify optimal reprogramming protocols, and predict how perturbations redirect developmental trajectories.

GRN constitutes the molecular foundation of differentiation, with TFs and their TGs forming intricate network that interpret signals and coordinate cellular responses. During differentiation, these networks undergo continuous rewiring that pluripotency factors are silenced while lineage-specific regulators activate in precisely timed cascades. Understanding these dynamic regulatory programs is therefore fundamental to simulating differentiation within AIVC.

Current computational approaches to GRN inference, however, fail to capture this essential dynamism. The prevailing methodology clusters cells into discrete types and infers static networks for each cluster, assuming regulatory relationships remain fixed within cell types. This static paradigm cannot explain the fundamental observation that apparently homogeneous cell populations give rise to heterogeneous fates. If regulatory networks were truly static within cell types, the mechanistic basis for divergent differentiation outcomes would remain unexplained. This limitation represents a critical barrier to implementing AIVC, as static models cannot simulate the continuous regulatory changes that drive differentiation. To address this challenge, we developed CellProphet, a framework that leverages temporal causality and transformer to infer dynamic GRN from single-cell transcriptomic data. By explicitly modeling how TF expression at earlier time points influences target gene expression at later stages, CellProphet captures the time-lagged dependencies inherent to gene regulation. The self-attention mechanism provides interpretable weights quantifying regulatory relationships, enabling both accurate prediction and biological insight.

We systematically validated CellProphet’s capabilities through a series of progressively complex experiments that demonstrate its ability to simulate cell differentiation within the AIVC framework. First, comprehensive benchmarking against nine competing methods across seven differentiation datasets established CellProphet’s foundational accuracy, achieving an average AUPRC of 0.541, a 162% improvement over the latest method CellOracle, confirming that temporal causality modeling captures regulatory relationships more accurately than conventional approaches. Building on this validated accuracy, we applied CellProphet to mouse embryonic stem cell differentiation where it identified 37 key TFs, including both well-established regulators like *Nanog, Pou5f1, Sox2* and potentially novel factors like *Nrf1, Klf4* and *Srebf1*. To verify that these identified TFs genuinely regulate their predicted targets, we integrated multi-modal epigenomic data including Hi-C chromatin loops, ATAC-seq accessibility, ChIP-seq binding, and histone modifications to confirm that CellProphet-inferred *Nanog* targets correspond to true regulatory interactions. Having established both TF identification and target validation, we demonstrated dynamic GRN reconstruction through Gaussian smoothing, revealing strong correlations (exceeding 0.7) between inferred regulatory strength and TF expression dynamics, confirming CellProphet captures genuine biological temporal patterns. These progressively validated capabilities, from basic accuracy to TF identification, target validation, and dynamic tracking, together contributed to successful *in silico* perturbation experiments where CellProphet accurately predicted cell fate transitions following *Gata1* and *Spi1* knockouts in hematopoietic differentiation, thereby achieving complete simulation of cellular differentiation responses and fulfilling the core promise of virtual experimentation within the AIVC framework.

The implications of CellProphet for future AIVC development are substantial. By providing dynamic GRN models that accurately predict perturbation outcomes, CellProphet enables virtual screening of thousands of genetic interventions to identify optimal therapeutic targets. The framework’s interpretability allows researchers to understand why specific perturbations produce particular outcomes, facilitating rational design rather than blind screening. As AIVC scales toward modeling entire organisms, CellProphet’s principle (temporal causality, interpretable architectures, and multi-scale flexibility) will remain foundational.

Several limitations warrant consideration for future development. First, CellProphet currently requires lineage-specific training, limiting generalization to novel systems. Foundation models trained on diverse datasets could enable zero-shot GRN inference. Second, the focus on transcriptional regulation, while capturing primary control mechanisms, omits chromatin accessibility, protein modifications, and metabolic regulation that also influence cell fate. Integration of multi-modal data will be essential for comprehensive virtual cells. Third, computational efficiency must improve to enable real-time simulation of complex perturbation combinations. Despite these challenges, CellProphet represents a critical advance toward realizing the AIVC vision, demonstrating that AI-driven approaches can capture differentiation dynamics with sufficient accuracy to transform how we understand and engineer cellular systems.

## Methods

### CellProphet algorithm overview

#### Theoretical Foundation

Our methodology is grounded on the core causal assumption that GRN is the intrinsic mechanism driving the dynamics of the transcriptome. We formally represent the time-series single-cell transcriptomic data as a matrix *E* ∈ ℝ^*T* × *G*^, where *T* is the number of cellular states ordered chronologically, and *G* is the number of genes. We posit that an accurate GRN matrix *A*∈ ℝ^*G* × *G*^ should encapsulate sufficient regulatory information to predict future cellular states based on historical expression data. This relationship can be formulated as:

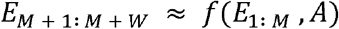

Here, E_1: *M*_ denotes the gene expression in watching window with length of *M*, while *E*_*M* + 1: *M* + *W*_ is the prediction of the future expression state in forecasting window with length of *W*. The function *f* models the complex, nonlinear kinetic relationships among genes. Consequently, we reframe the problem of GRN inference as an optimization task: to find the best regulatory matrix *A*^*^ that minimizes the discrepancy between the model’s predictions and the actual observations:

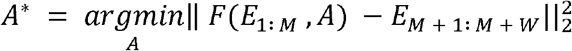

#### Time series expression data construction

In each of the seven datasets provided by BEELINE, our raw data is a gene expression matrix *E* ∈ ℝ ^*c* × *G*^, where *C* and *G* represent the number of cells and genes, *E*_*ij*_ indicates the expression value of gene *j* within cell *i*. To endow the raw data with temporal information, we employ the Slingshot algorithm to calculate a pseudotime for each cell. Then we sort the cells in chronological order based on the pseudotime, thus transforming the raw expression data into a time series data *E* ∈ ℝ ^*T* × *G*^, where *T* is the number of time points. It’s worth noting that time points *T* is equal to cell numbers *C*, as each cell corresponds to a pseudotime on differentiation trajectory. In this way, our time series data *E =* [*e_1_,e_2_*,…,*e*_*T*_]^*T*^ possesses the temporal relationships of genes during the differentiation process as we map each cell to a pseudotime, where *e*_*i*_ ∈ ℝ ^*G*^ represents the gene expression at the *i*-th time point.

After that, we split time series data *E* into dataset *D* ={(*X*_*i*_,*Y*_*i*_) | *i* =1,2,…,*N*}, where *X*_*i*_ are input samples, *Y*_*i*_ are the ground truth associated to each sample and *N* is the number of samples. Each sample *X*_*i*_ =[*e*_*i*_,*e*_2_,…,*e*_*i*+*M*-1_]_*T*_ ∈ ℝ ^*M* × *G*^ and *Y*_*i*_ [*e*_*i* + *M*_,…,*e*_*i*+*M*+*W*-1_]^*T*^ ∈ ℝ^*W* × *G*^ is the gene expression in watching window of length *M* and forecasting window of length *W* respectively. Therefore, CellProphet takes the gene expression in watching window as input and predicts the expression in forecasting window, ultimately obtaining the final GRN.

#### Model architecture

CellProphet leverages a Transformer-based architecture to infer causal gene regulatory networks from time-series expression data. It is composed of two primary components: The Temporal Aggregation module for integrating dynamic temporal patterns and the Regulatory Filtering module for refining putative causal interactions. Initially, the input data *X* ∈ ℝ ^*M* × *G*^ is processed by a fully connected layer to embed each expression value, yielding an embedding matrix 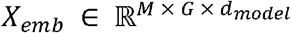, where *d*_*model*_ is the embedding dimension. Subsequently, this matrix is transposed to 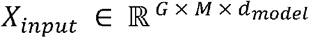, which will be input to Temporal Aggregation module. The core function of the Temporal Aggregation module is to construct a condensed, context-aware representation of historical gene expression. For a given time point *t*, to predict the expression dynamics over future states (*t* + 1, *t* + 2,…, *t* + *w*), the model must consider the entire history of expression states for all genes up to time *t* (i.e., from time points 1,2,…,*t* − 1). The core principle of this module is that all previous stages count, not just a single time point and previous time points contribute unequally to the prediction of future gene expression, so we need to assign differential weights to individual time point. For instance, if a gene maintains a stable expression level before being suddenly up-regulated at a specific time point *m*, this event at *m* is likely to be more determinant of future expression and should thus be assigned a greater weight than other less eventful moments. To computationally implement this principle, the module employs the self-attention mechanism inherent to the Transformer architecture to learn the relative importance of different historical time points. This mechanism computes attention weights that quantify the significance of each past expression state for future prediction. Specifically, we introduce a temporal mask *Φ*^*t*^ ∈ ℝ^*M* × *M*^ in the form of a lower-triangular matrix:

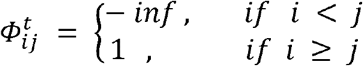

In this scheme, an entry 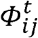, representing the attention weight between time point *i* and time point *j*, is computed (i.e., is non-zero) if and only if *i* ≥ *j*. This trick is in line with the principles of cell differentiation, as a cell at a specific time can only know the gene information from its ancestors but not its descendants, because they haven’t been differentiated yet at this time point. Thus, the calculation in Temporal Aggregation module is modified as follows:

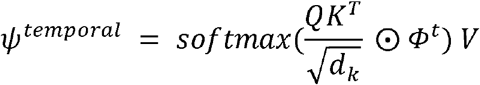

where Query (*Q*), Key (*K*), Value (*V*) matrices are generated by linear transformation of the input embedding 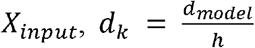 (*h* is the number of heads in multi-head self-attention) and represents hadamard product. 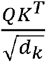 yields an attention matrix *A* ∈ ℝ^*G* × *M* × *M*^, *A*_*ijk*_ represents the attention score of gene *i* between the time point *j* and time point *k*. The softmax operation then normalizes the attention scores between different time points into attention weights that sum to one, which represent the relative importance of the association between these time points for a given gene. Finally, a context-aware representation of each gene is generated by taking the weighted sum of the Value (*V*) matrix, where the weights are the output of softmax operation. This step effectively aggregates information from different time points based on their learned importance. The formula for this final weighted sum is:

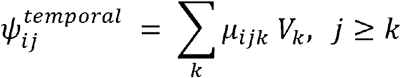

where 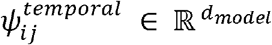 represents the feature for gene *i* after aggregating historical expression information before time point *j, μ*_*ijk*_ represents the attention weight for gene *i* between time point *j* and *k* (where *j* ≥ *k*) and 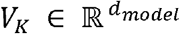 represents the expression feature of gene *i* at the historical time point *k*. The output of Temporal Aggregation module 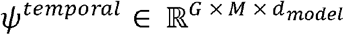 will be transposed to 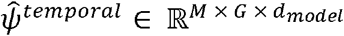, which will be input into Regulatory Filtering module alongside the prior information to learn the regulatory coefficient between genes. The prior information we employ is a comprehensive database NicheNet, which itself integrates data from over 50 public sources about gene regulations for human and mouse. In addition, we refine the prior information by merging its edges with those from the ground truth GRN, ensuring that the edges to be inferred are included within the prior. Since the prior can be represented as a graph, we use its adjacency matrix *P ∈* ℝ^*G* × *G*^ as the regulatory filter mask:

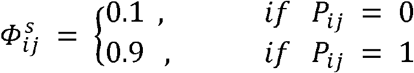

*Φ*^*s*^ assign a higher initial weight to gene pairs with interactions documented in prior, while all other potential pairs receive a small, non-zero baseline weight. We experiment with multiple combinations for weight allocation and ultimately assign a weight ratio of 1:9. The weight allocation strategy encourages the model to prioritize the refinement of known regulatory pathways while still allows for the discovery of novel links (via the non-zero weight), we effectively balance established biological priors with data-driven, de novo discovery. Thus, the calculation in Regulatory Filtering module can be formulated as follows:

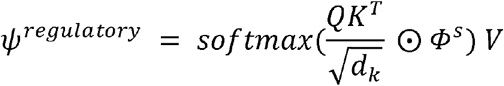

where Query (*Q*), Key (*K*), Value (*V*) matrices are generated as linear transformation of 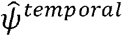. Ultimately, the output, *ψ*^regulatory^ will go through a linear layer to obtain the predicted gene expression *Ŷ*_*i*_ ∈ ℝ^*W* × *G*^. We use MSE loss as our loss function:

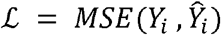

### Cell lineage GRN inference

During inference, the sampling strategy for the test data differs from that used in the training phase, as outlined below:

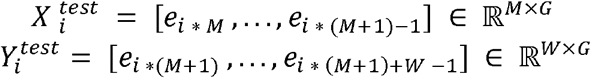

The time series data are divided into segments, each with a length of *M* corresponding to the watching window. These segments are processed through the trained CellProphet, each watching window after being processed will result in an attention matrix *A* ∈ ℝ ^*M* × *G* × *G*^ in Regulatory Filtering module. Assuming the number of cells in the lineage is *C*, a total of 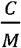 attention matrix will be generated (for simplicity, we assume that *C* is divisible by *M*). The generated attention matrices are then averaged to produce a single, aggregate attention matrix *A*^*^ ∈ ℝ ^*G* × *G*^. For gene *i*, we use the absolute alues of log2 fold changes to measure its differential expression levels between the start and all the subsequent states and then take their average across all the developmental states, denoted as 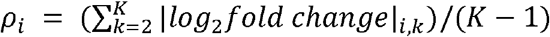.Here, the developmental states are simply annotated by equally dividing cells into *K* parts along the pseudotime trajectory (*K* =4 in our paper). For each TF-TG edges, we compute their regulatory score:

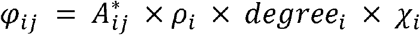

where 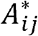 is the attention weight between gene *i* and gene *j, degree*_*i*_ refers to the number of genes targeted by the gene *i* and *χ* _*i*_ indicates the number of target genes with *ρ* > 1 for gene *i*. We assume that a TF which has a greater number of targeted genes contributes more to the network. Additionally, if the target genes have *ρ* > 1, indicating that these genes show significant changes during development and are likely contributors to the developmental process. Thus, multiplying by *χ* _*i*_ acts as a reward term to the gene *i*. We finally select the top *k* edges based on the regulatory score *φ* _*ij*_ to construct the cell lineage GRN, where *k* corresponds to the number of edges in the ground truth GRN.

#### Dynamic GRN inference

During the inference stage, testing data are sampled based on watching windows. Given a lineage containing *C* cells, a total of 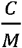 watching windows can be generated, where *M* denotes the length of the watching window. Each window, after being processed by the trained CellProphet, generates a GRN. These GRNs each correspond to distinct time periods along the developmental trajectory. The averaging step in lineage GRN inference is omitted. Instead, the 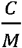 inferred GRNs are temporally smoothed by Gaussian smoothing. Specifically, for a specific TF-TG pair, we can obtain their 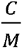 regulatory scores at different developmental time periods. Gaussian smoothing computes a new, smoothed regulatory score *Ŝ*_*j*_ for each time period *j* by taking a weighted average of all points in the original regulatory score. The weights are determined by a Gaussian kernel. For each time period *j* we are calculating, the weight assigned to any other time period *i* is given by:

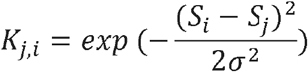

where *S*_*i*_ is the original regulatory score at period *i*, and the standard deviation *σ* controls the width of the smoothing window. A larger *σ* results in a smoother curve. In our implementation, *σ* is systematically derived from a hyperparameter Full Width at Half Maximum (FWHM):

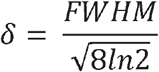

In our paper, we set FWHM to be 8. Before applying the weights, the kernel is normalized to ensure the weights sum to 1, thus preserving the overall magnitude of the signal. The normalized weight is 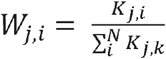, where *N* is the total number of time periods (i.e., 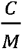). The final smoothed regulatory score *Ŝ*_*j*_ for time period *j* is the convolution of the normalized kernel with the original regulatory score *S* :

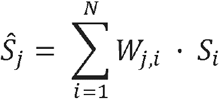

This process effectively filters out high-frequency fluctuations, allowing for a clearer, more robust analysis of the principal dynamic regulatory patterns between a TF and its target genes.

#### Training details

For the model architecture, we set the lengths of both the watching window and forecasting window to 16. The model was composed of 4 Transformer blocks, with a model dimension *d*_*model*_ of 128 and 4 attention heads within each block. For optimization, we employed the Adam optimizer. The training process began with a warmup strategy that linearly increased the learning rate from 0 to 1e-4, followed by a CosineAnnealingLR scheduler for subsequent adjustments. The model was trained for a maximum of 20 epochs on an 80G Nvidia A100 GPU. To prevent overfitting, an early stopping strategy was implemented with a patience of 3 epochs. With a batch size of 4, the training process for each dataset took approximately 30 minutes and occupied 40 GB of GPU memory.

#### Prior information NicheNet

Our study utilized large-scale, directed graphs as prior networks for both human and mouse gene regulation. The primary prior was adapted from the NicheNet resource, a comprehensive database integrating over 50 public data sources. We first tailored this network to our needs through several processing steps. Specifically, we focused on intracellular processes by filtering out ligand-receptor interactions, used the unweighted version of the network, and ensured all connections were directed by treating undirected edges as bidirectional. To generate a version for mouse studies, the human gene symbols from the processed network were mapped to their one-to-one mouse orthologs using ENSEMBL^39^, genes lacking a clear ortholog were excluded. The final curated networks comprised 25,332 genes and 5,290,993 edges for human, and 18,579 genes and 5,029,532 edges for mouse.

### Datasets

#### mESC dataset^40^

This dataset comprises scRNA-seq data for 421 primitive endoderm differentiated from mouse embryonic stem cells, collected across five time points: 0, 12, 24, 48, and 72 hours. BEELINE utilized Slingshot to calculate pseudotime for each cell, with cells measured at 0h as the starting cluster and the cells measured at 72h as the ending cluster.

#### mHSC dataset^41^

This dataset consists of scRNA-seq data from 1,656 hematopoietic stem cells undergoing differentiation. Notably, the dataset is divided into three lineage-specific datasets: mHSC-E (Erythroid lineage), mHSC-GM (Granulocyte-Monocyte lineage), and mHSC-L (Lymphoid lineage). BEELINE applied Slingshot to independently calculate pseudotime for cells within each of the three lineages.

#### mDC dataset^42^

This dataset contains scRNA-seq data from over 1,700 mouse bone-marrow-derived dendritic cells stimulated with lipopolysaccharide and measured at 1, 2, 4, and 6 hours. BEELINE computed the pseudotime using Slingshot, with cells measured at 1h as the starting cluster and cells measured at 6h as the ending cluster.

#### hESC dataset^43^

This dataset consists of a time-course scRNA-seq experiment with 758 human embryonic stem cells differentiating into definitive endoderm cells, measured at 0, 12, 24, 36, 72, and 96 hours. BEELINE computed the pseudotime using Slingshot, with cells at 0h as the starting cluster and cells at 96h as the ending cluster.

#### hHep dataset^44^

This dataset is from an scRNA-seq experiment tracking the differentiation of induced pluripotent stem cells into hepatocyte-like cells. It includes 425 cells measured at multiple time points: days 0, 6, 8, 14, and 21. BEELINE calculated the pseudotime using Slingshot, setting cells from day 0 as the starting cluster and cells from day 21 as the ending cluster.

### Cell lineage GRN benchmarking

BEELINE provided the ground truth GRN for each dataset obtained from ChIP-seq data. Assuming the total number of genes is *G*,the total number of potential edges is *G* × − 1), where the subtraction of 1 excludes self-loop edges. Among these edges, those present in the ground truth GRN are labeled as 1 (positive edges), while the remaining edges are labeled as 0 (negative edges). Thus, the evaluation of GRN can be treated as a binary classification problem, where the task is to determine whether the edges identified by the model exist in the ground truth GRN. We computed AUROC, AUPRC, F1 score and EPR as metrics. The PR curve describes the relationship between recall and precision, while the ROC curve illustrates the relationship between false positive rate (FPR) and true positive rate (TPR). Their definitions are as follows:

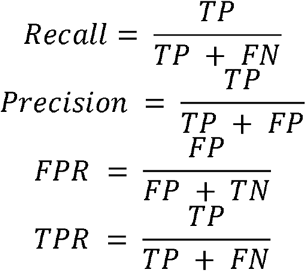

where TP (True Positive) refers to edges predicted as true by the model that are also present in the ground truth GRN. FP (False Positive) refers to edges predicted as true by the model but absent from the ground truth GRN. TN (True Negative) refers to edges predicted as false by the model that are also absent from the ground truth GRN, while FN (False Negative) refers to edges predicted as false by the model but present in the ground truth GRN. Subsequently, the calculations of AUROC, AUPRC, and F1 score are as follows:

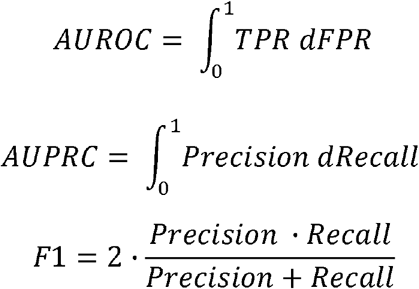

As for EPR, BEELINE first defined a term *early precision* as the fraction of true positives in the top *k* edges. Thus, EPR represents the ratio of *early precision* value calculated by algorithm and the *early precision* calculated by a random predictor. Specifically, the random predictor’s precision is the edge density of the ground truth network.

### AUCell algorithm

To identify key TFs during cellular differentiation based on the inferred GRN, we employed the AUCell algorithm to calculate the activity score for each TF. AUCell first ranked gene expression in each cell from high to low and set a threshold (defaulting to the top 5%) to identify highly expressed genes in that cell. For each threshold (ranging from 0 to 5%), the algorithm calculated the proportion of genes within the TF regulon that are highly expressed in the given cell. By computing the area under the curve (AUC), the algorithm determined the activity score of each TF in the cell. A higher score indicated that the TF is more actively expressed in the given cell. Since the mESC dataset included five sequencing time points (0h, 12h, 24h, 48h, and 72h), we performed ANOVA on the TF activity scores across these time points. TFs with a p-value less than 0.01 were identified as key regulators in the differentiation process.

### mESC Hi-C and scATAC-seq data processing

To identify the TF-TG edges for constructing dynamic GRN, we utilized previously published experimental dataset^30^ to validate the accuracy of the edges inferred by CellProphet. The mouse Hi-C and scATAC-seq data from the dataset were processed and mapped into mouse reference genome (mm9 version, UCSC). For Hi-C data, raw FASTQ files were processed using HiC-Pro^45^ to generate valid pairs and the contact matrix. To convert the valid pairs into “.hic” format, we used the “pre” function from JuicerTools, followed by chromatin loops identification at a 20 kb resolution using the “hiccups” function. For scATAC-seq data, reads were mapped to the mm9 reference genome using Bowtie2^46^, with mitochondrial reads and those overlapping blacklist regions removed. Following prior studies^47,48^, to account for Tn5 transposase binding as a dimer and inserting adapters separated by 9 bp, we adjusted the read positions: reads aligned to the (+) strand were shifted by +4 bp, while those aligned to the (-) strand were shifted by −5 bp. Peak calling was performed using MACS^49^, and TF footprints were identified using the “footprinting” function in Regulatory Genomics Toolbox (RGT)^50^. To analyze *Nanog* regulatory regions, we retrieved the *Nanog* motif from JASPAR^51^ and scanned the identified footprint regions to locate *Nanog* footprint.

Finally, we integrated the chromatin loops derived from Hi-C data, histone modification data (H3K9me3, H3K27me3, H3K4me1, H3K4me3, and H3K27ac), peaks identified from scATAC-seq data, *Nanog* footprint, and *Nanog* ChIP-seq data obtained from ChIP-Atlas^33^. These data were visualized collectively using IGV, enabling the comprehensive evaluation of the regulatory landscape.

### *In silico* perturbation

#### Perturbation vector calculation

We followed CellOracle’s *in silico* perturbation implementation to validate whether CellProphet-inferred lineage GRN could simulate cell fate changes after TF knockout. We set the perturbed TF’s expression to zero, generating an expression difference vector relative to its original value. This vector was multiplied by CellProphet ‘s GRN coefficient matrix, the average of matrices from 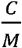 watching windows, where *C* represented lineage cell count and *M* denoted watching window length. The GRN coefficient matrix encoded the regulatory strength between each TF and its TG. For each TF-TG pair, we computed the PCC of their expression profiles. When PCC < 0, we inverted the sign of the corresponding coefficient to reflect repressive regulation. During propagation, we reset the perturbed gene’s expression change to its original perturbation value after each iteration, ensuring persistent effects. Following propagation, each cell’s final expression difference vector was used to calculate transition probabilities:

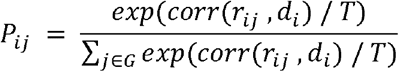

where *d*_*i*_ was the final expression difference vector for cell *i, r i*_*j*_ was the origin gene expression difference between cell *i* and cell *j*, corr operation denoted Pearson correlation, and *T* was the temperature parameter. These probabilities governed transitions from cell *i* to cell *j*. We then computed the perturbation vector for cell *i* by weighting diffusion-map^52,53^ coordinate differences (*V*_*ij*_) with transition probabilities:

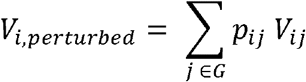

This vector indicated the direction of phenotypic change induced by perturbation.

#### Perturbation score calculation

We calculated the gradient of the pseudotime using the “numpy.gradient” function, resulting in a two-dimensional vector representing the differentiation direction. Subsequently, we computed the inner product between the pseudotime gradient vector and the perturbation vector, which was used as the perturbation score (PS). A positive PS indicated that the TF knockout promoted the current cell’s development, as its differentiation direction aligned with the perturbation. Conversely, a negative PS signified inhibition, indicating opposing directions.

### Gene expression prediction after perturbation

In addition to predicting cell fate changes after perturbation based on the perturbation vector, we collected gene expression data associated with *Gata1* knockout from Gene Expression Omnibus (GEO) (GSE279985) to evaluate CellProphet’s ability to predict expression changes following perturbation. This dataset includes gene expression from three wild-type and three knockout mice. We averaged expression levels for each condition and computed the per-gene difference as the ground truth. We then utilized the final expression difference vector generated by the *in silico* perturbation as the predicted gene expression profile following *Gata1* perturbation and evaluated whether the predicted direction of gene expression changes aligned with the ground truth.

### Statistical Analysis

For the pairwise statistical comparisons with 9 benchmark methods across all datasets, we employed the Mann-Whitney U test. For the identification of key TFs in mESC differentiation, after calculating the activity score for each TF regulon, we partitioned cells into five time intervals based on mESC sampling time and performed ANOVA analyses within these intervals to compute the p-value for each TF regulon. To assess whether changes in regulatory strength were consistent with TF expression dynamics, we calculated the PCC and R-squared between the actual expression of each TF and the regulatory scores of its TF-TG edges.

## Acknowledgments

This research was supported by National Key Research and Development Program of China (2024YFF0507400), National Natural Science Foundation of China (6220071694), and the Macao Young Scholars Program (AM2023024).

## Author Contributions

Conceptualization: P.R. and J.W. Data acquisition: P.R. and J.W. Analysis: P.R. and J.W. Methodology: P.R., J.W. Supervision: J.W. and Y.G. Funding acquisition: J.W. and Y.G. Project administration: J.W. Writing: P.R., Y.G., Y.C., and J.W.

## Competing Interests statement

The authors declare no competing financial interests.

## Data availability

All the datasets analyzed in this study are publicly available. The scRNA-seq datasets are available in the GEO under accession codes: GSE98664[https://www.ncbi.nlm.nih.gov/geo/query/acc.cgi?acc=GSE98664] (mESC), and GSE81682[https://www.ncbi.nlm.nih.gov/geo/query/acc.cgi?acc=GSE81682] (mHSC). The prior gene interaction network was from NicheNet, which can be downloaded from https://github.com/saeyslab/nichenetr. The ChIP-seq data for validating the constructed GRN were obtained from BEELINE. Epigenomic datasets for mESC, including scATAC-seq, histone modification, and Hi-C data, are available in the GEO under accession codes: GSE159623[https://www.ncbi.nlm.nih.gov/geo/query/acc.cgi?acc=GSE159623]. The *Gata1* knockout data for mHSC can be accessed in the GEO under accession codes: GSE279985[https://www.ncbi.nlm.nih.gov/geo/query/acc.cgi?acc=GSE279985]. Source data are provided with this paper.

## Code availability

The CellProphet algorithm is implemented in Python. The source code of CellProphet is available at https://github.com/prsigma/CellProphet.

---

**Extended data Figure 1.**
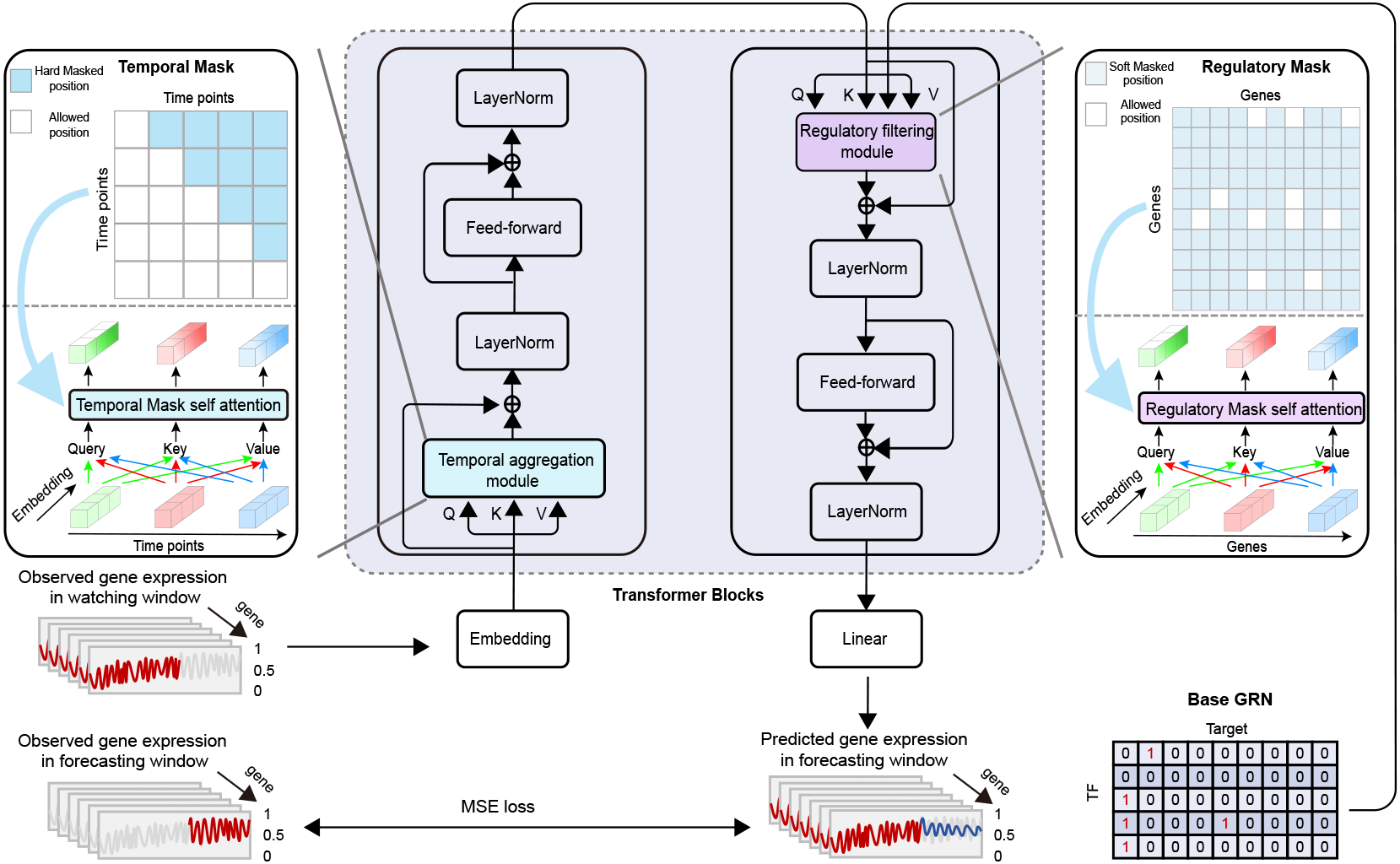
CellProphet architecture. CellProphet is built on Transformer encoders, where Temporal and Regulatory masks are specifically designed to compute self-attention. The model takes observed gene expression as input to predict gene expression over a specified forecasting window. The attention map produced by the Regulatory Filtering module is interpreted as the GRN coefficient matrix. CellProphet infers an optimal GRN by minimizing the discrepancy between predicted and actual gene expression.

**Extended data Figure 2.**
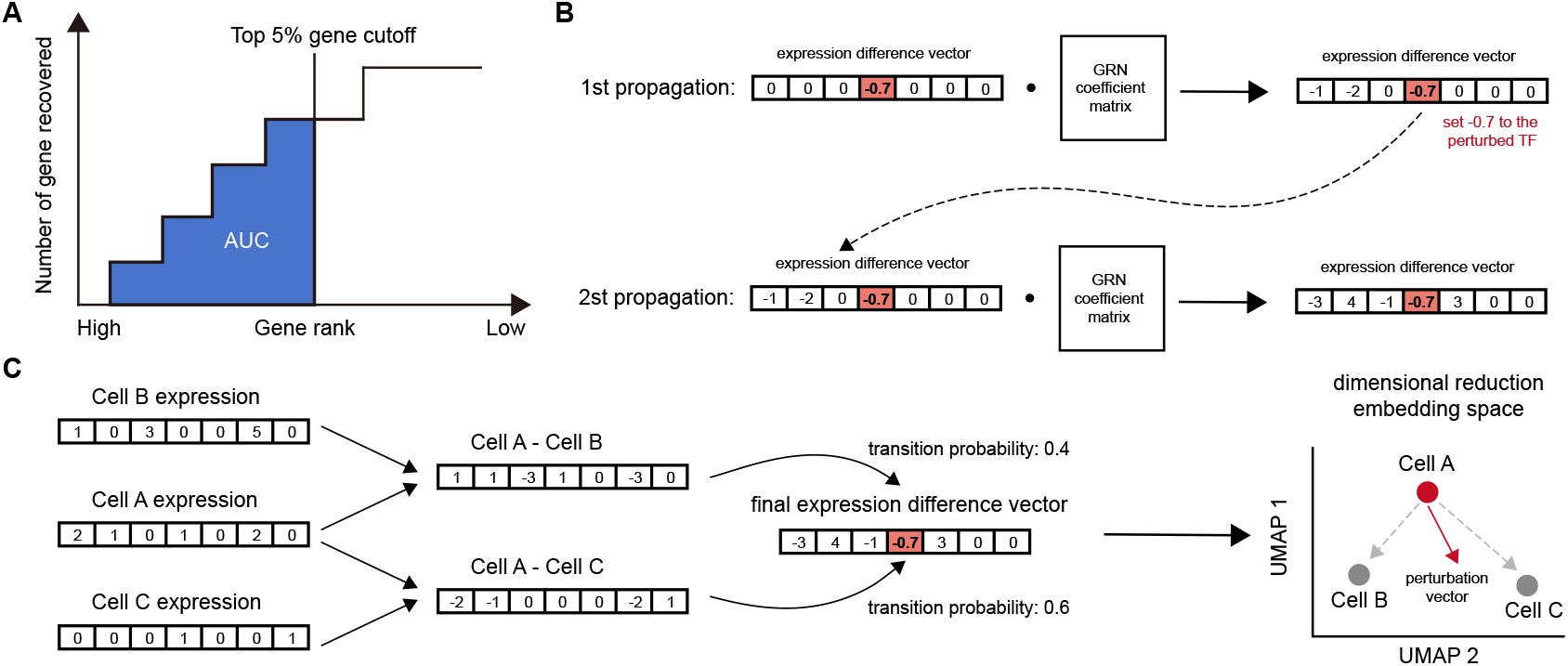
GRN inference process and workflows of AUCell, *in silico* perturbation. **A**, Threshold of AUCell is progressively increased to determine how many genes in each TF regulon are highly expressed in a given cell. The area under the resulting curve is computed as the TF’s activity score. **B**, The effects of gene perturbations are propagated according to the GRN coefficient matrix. After each propagation step, the value of the perturbed gene is reset to its initial perturbation value to ensure the perturbation’s influence persists throughout the whole propagation process. **C**, Once the final expression difference vector for a cell is obtained, the transition probabilities are calculated by comparing the vector to the original expression difference vectors of its neighboring cells. These probabilities are combined with the coordinate difference vectors in the 2D space of the cells to compute a weighted sum, resulting in the perturbation vector for the cell.

**Extended data Figure 3.**
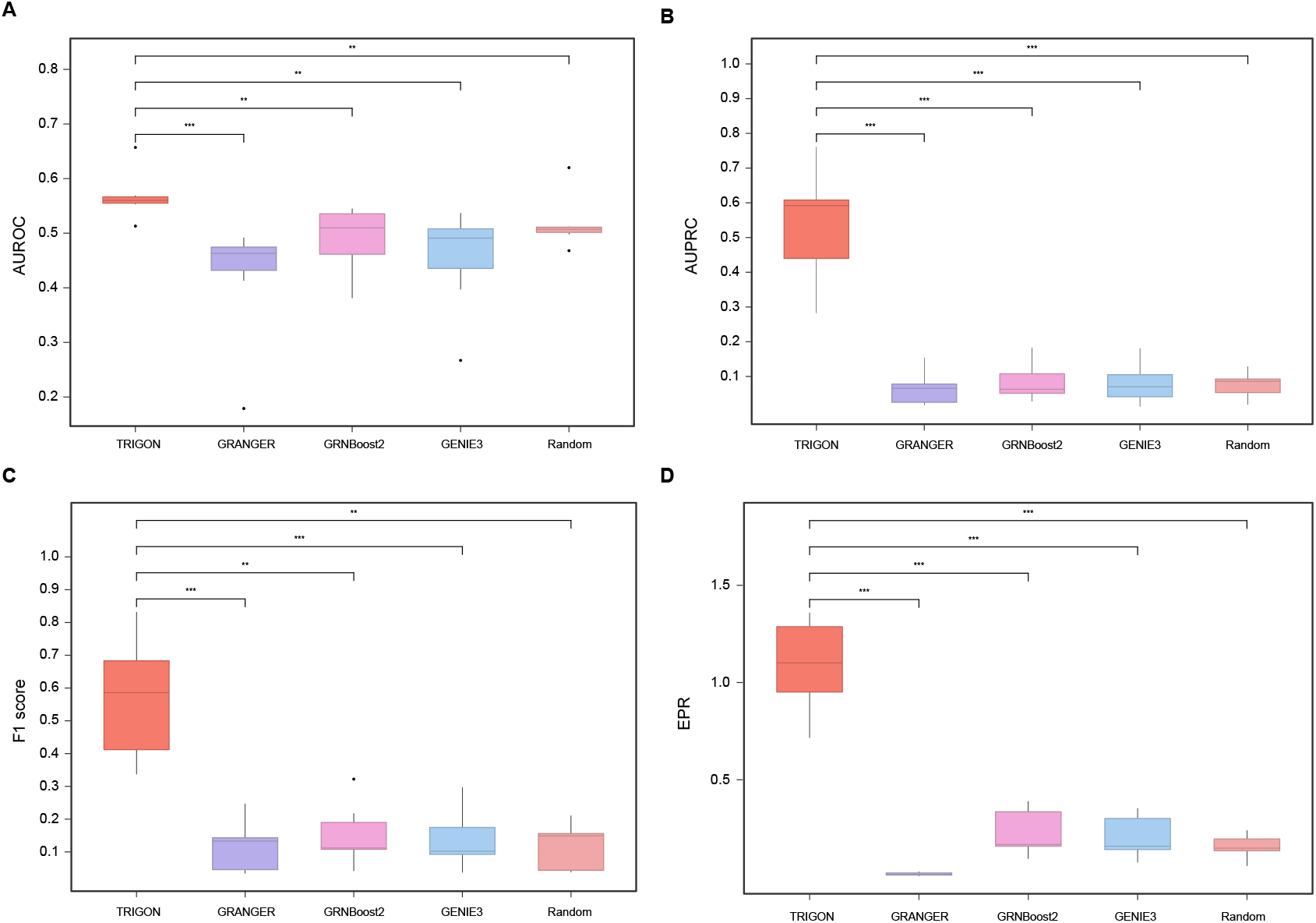
Performance of methods without prior information. Box plots comparing AUROC **(A)**, AUPRC **(B)**, F1 score **(C)**, and EPR **(D)** across seven datasets for different methods excluding the use of prior information. Statistical significance was assessed using the Mann-Whitney U test. \**P*<0.05, \*\**P*<0.01, \*\*\**P*<0.001.

**Extended data Figure 4.**
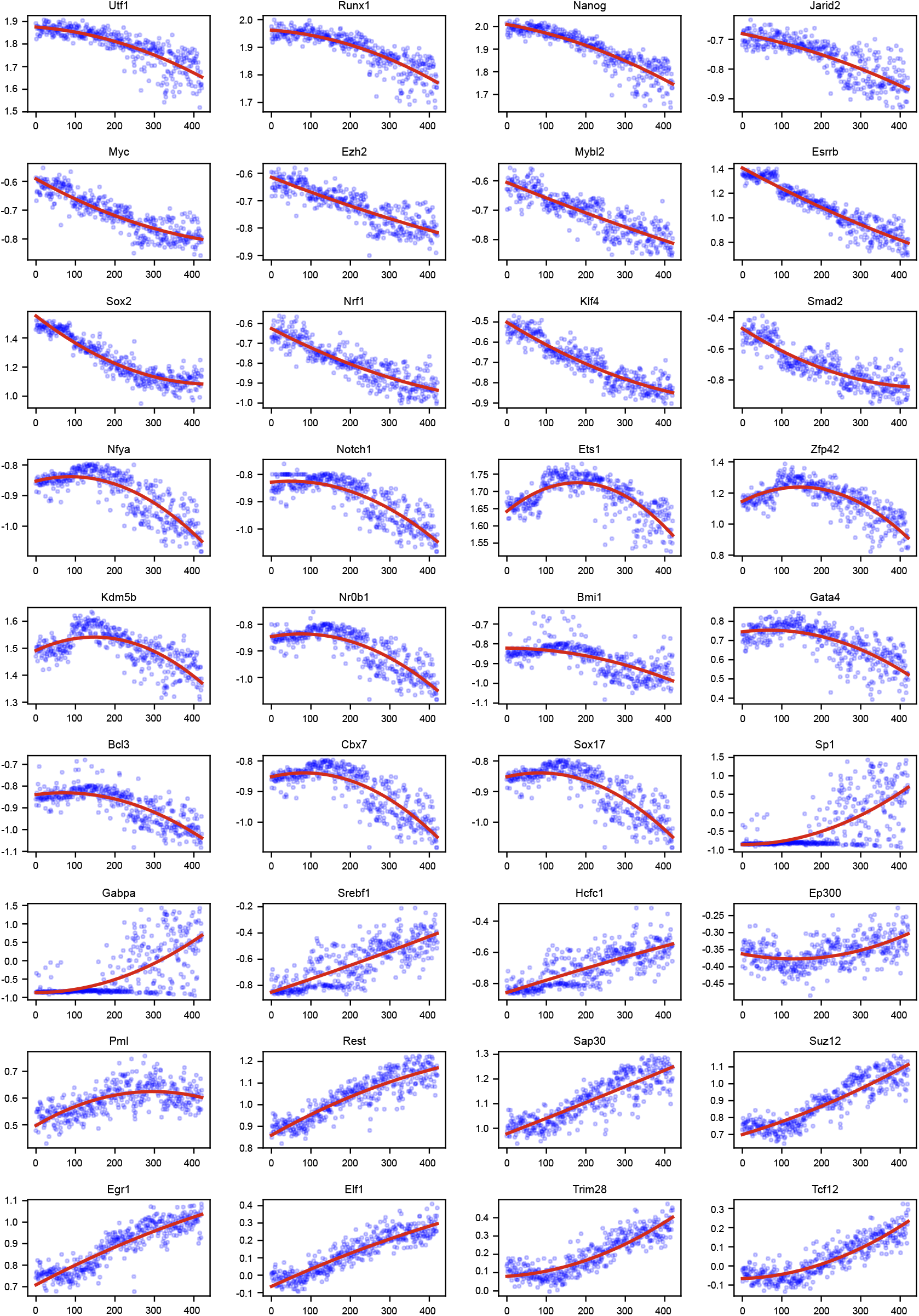
The activity score trends of key TFs identified by CellProphet. These activity scores were fitted with quadratic curves, revealing three distinct patterns of dynamic change.

**Extended data Figure 5.**
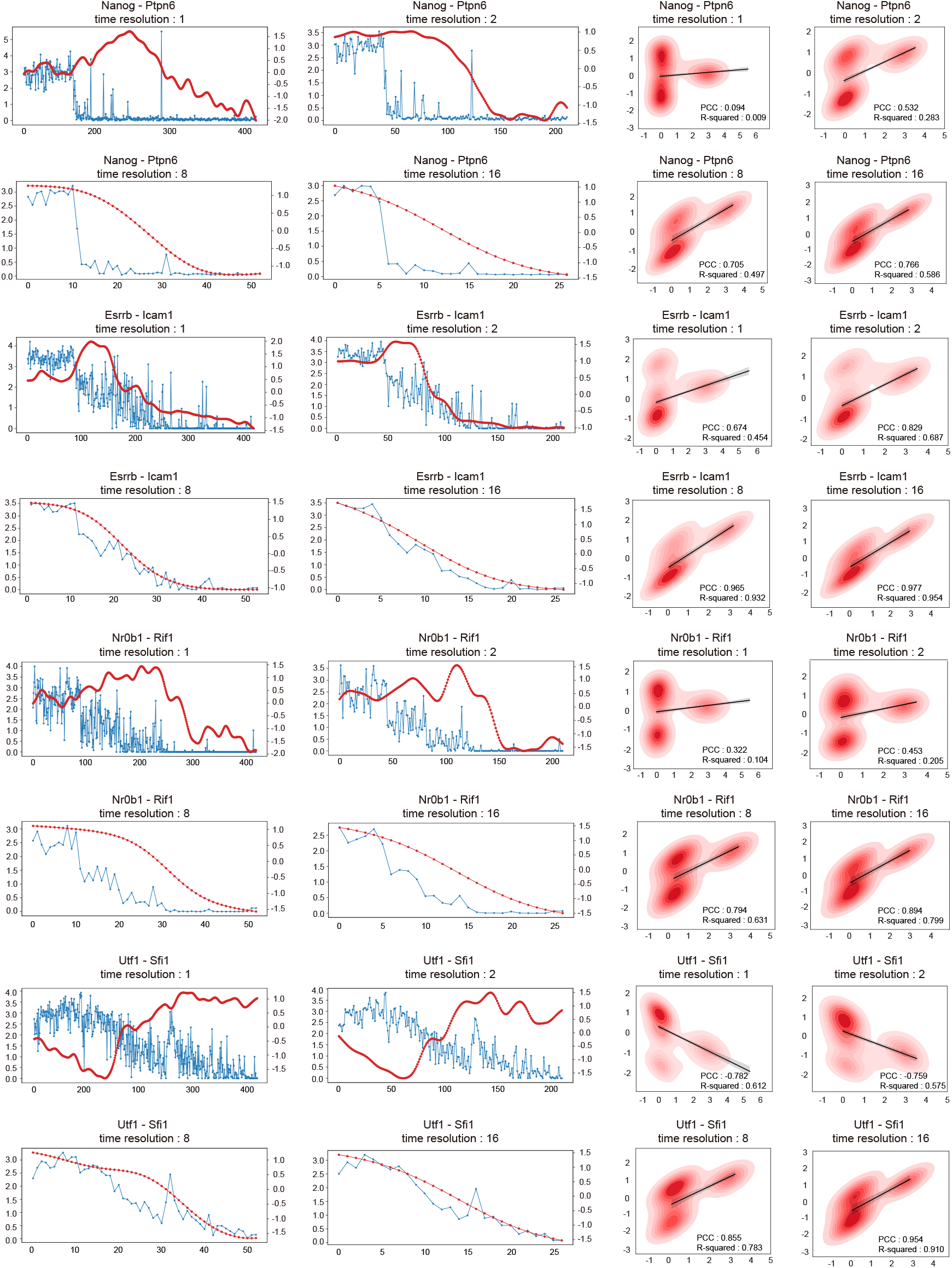
CellProphet constructs dynamic GRN at different temporal resolutions. At resolution of 1, the PCC between the regulatory strengths and the corresponding gene expression is the lowest. As the resolution changes to 16, the PCC steadily improve, reaching above 0.7.

**Extended data Figure 6.**
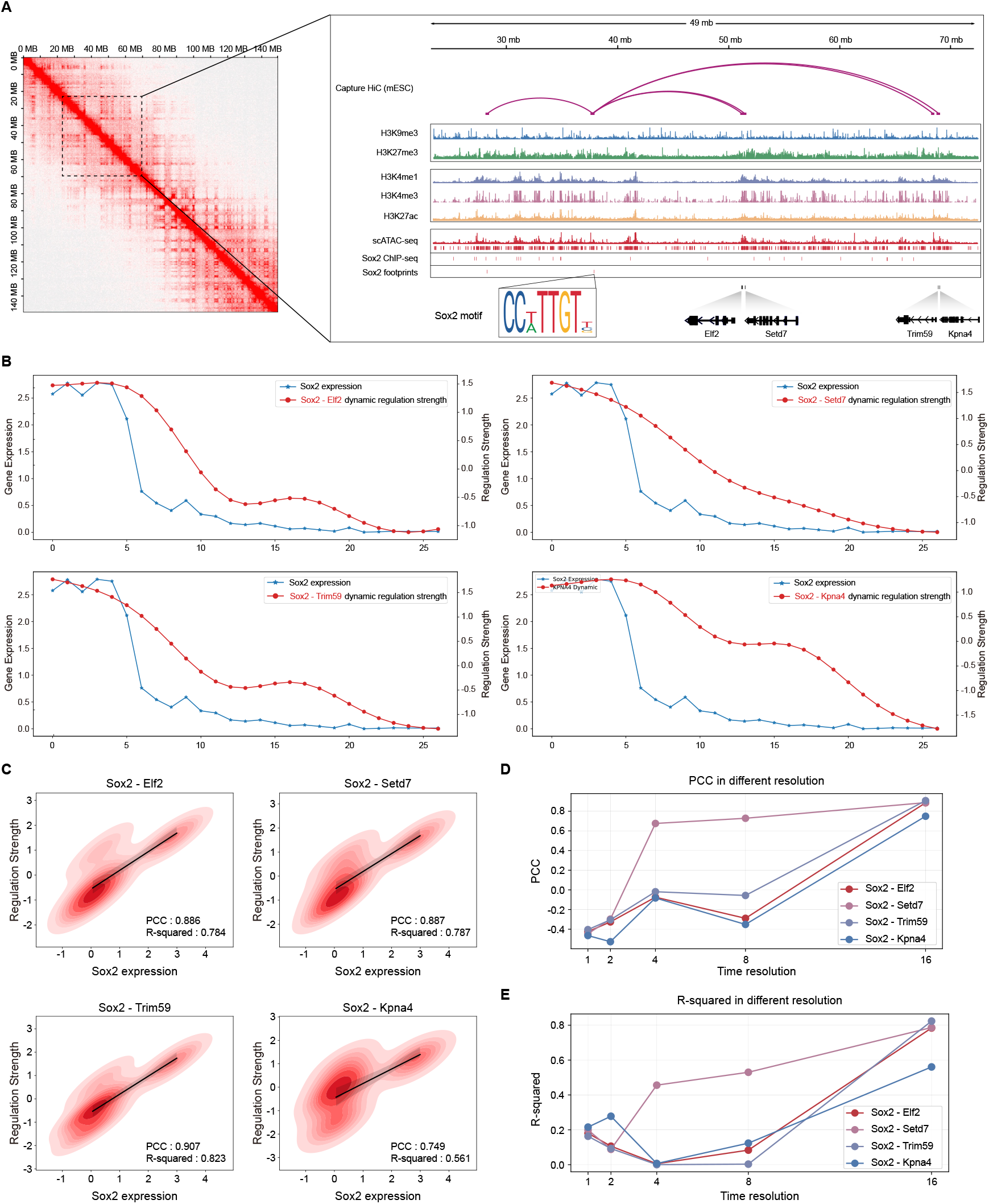
The dynamic GRN of *Sox2*. **A**, Four target genes of *Sox2* (*Elf2, Setd7, Trim59, Kpna4*) inferred by CellProphet are validated by Hi-C and scATAC-seq data. **B**, The dynamic GRN of *Sox2* at temporal resolution of 16. **C**, In the inferred *Sox2* dynamic GRN, regulatory strength shows a high correlation with *Sox2* gene expression. **D-E**, Comparison of PCC and R-squared for the *Sox2* dynamic GRN across different temporal resolutions.

**Extended data Table 1.** Label distribution. Across the seven datasets, positive edges account for only 0.9%, 0.65%, 0.7%, 1.15%, 0.22%, 0.17% and 0.24% of all potential edges, respectively. This indicates that positive labels are significantly outnumbered by negative labels, resulting in severe class imbalance.

|  | mESC | mHSC-E | mHSC-GM | mHSC-L | mDC | hHep | hESC |
| --- | --- | --- | --- | --- | --- | --- | --- |
| Genes | 1652 | 1933 | 1520 | 640 | 599 | 1638 | 1747 |
| Potential edges | 2727452 | 3734556 | 230880 | 408960 | 358202 | 2681406 | 3050262 |
| Positive edges | 24557 | 24726 | 16198 | 4705 | 796 | 4476 | 7411 |
| Proportion of positive edges | 0.90% | 0.65% | 0.70% | 1.15% | 0.22% | 0.17% | 0.24% |

**Extended data Table 2.**
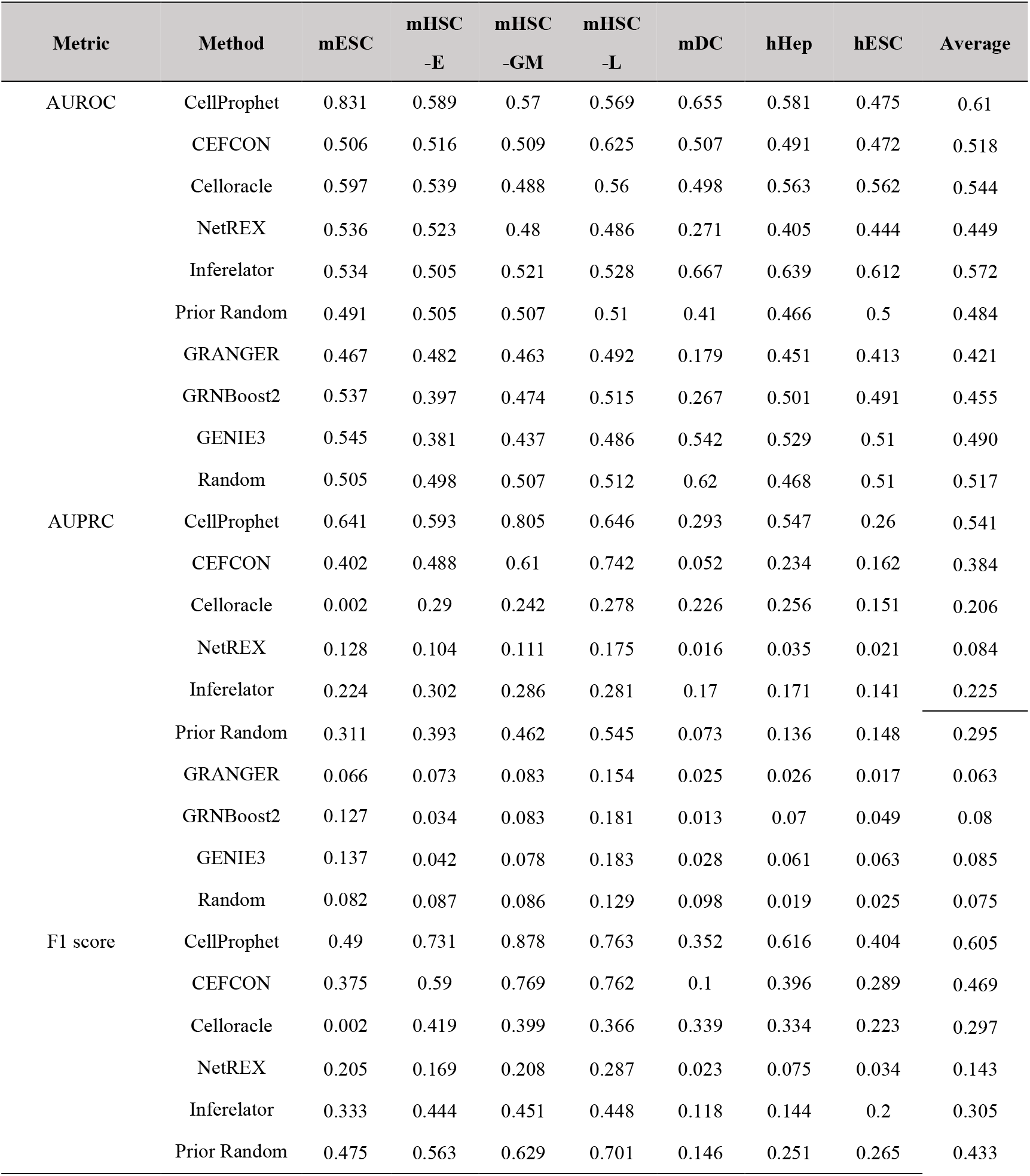
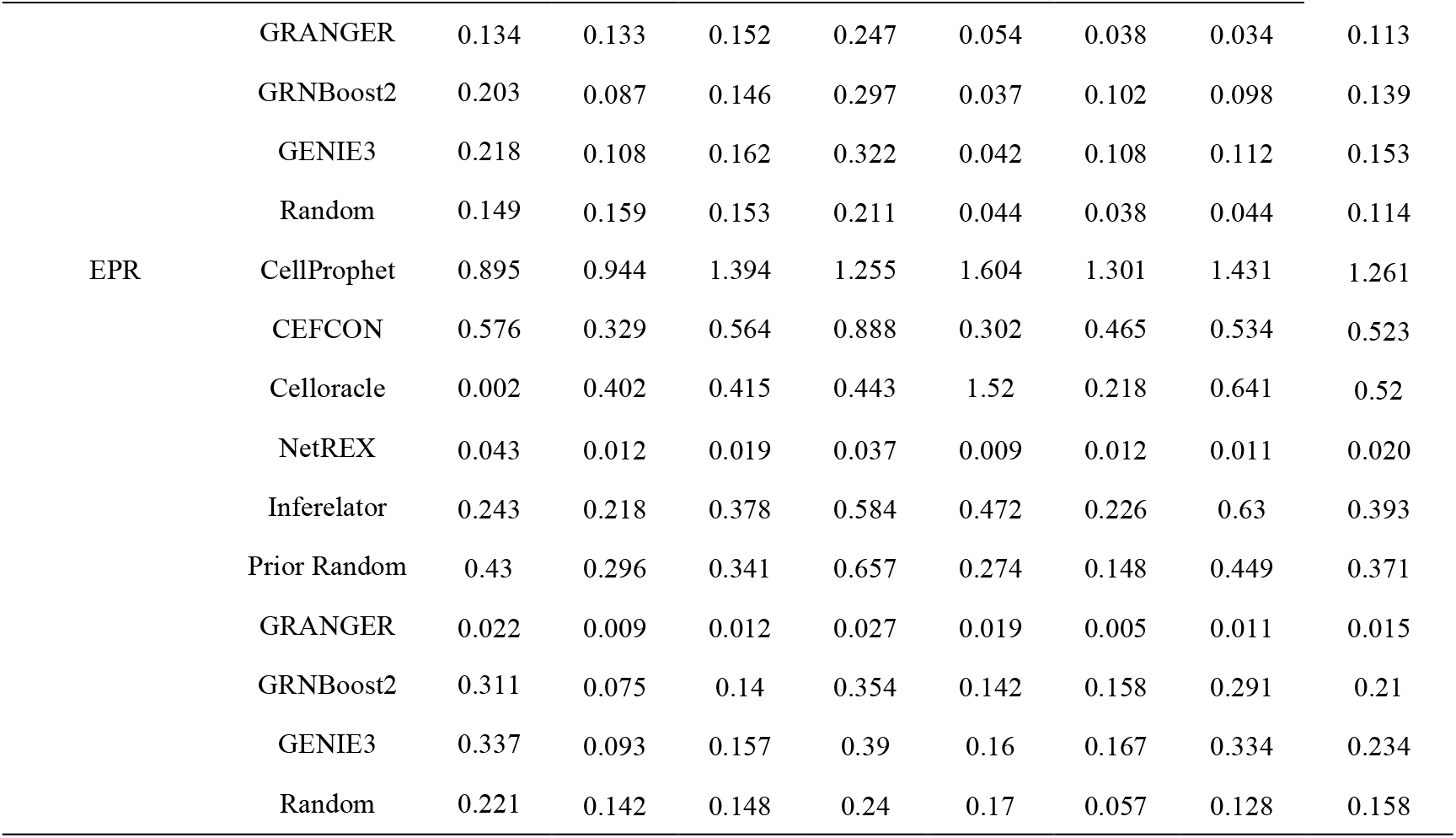
Performance comparison of GRN inference methods using the prior information.

**Extended data Table 3.** Performance comparison of GRN inference methods without prior information.

**Extended Table 3.** Performance comparison of GRN inference methods without prior information.
| Metric | Methods | mESC | mHSC<br>-E | mHSC<br>-GM | mHSC<br>-L | mDC | hHep | hESC | Average |
| --- | --- | --- | --- | --- | --- | --- | --- | --- | --- |
| AUROC | CellProphet | 0.657 | 0.564 | 0.553 | 0.557 | 0.56 | 0.569 | 0.513 | 0.568 |
|  | GRANGER | 0.467 | 0.482 | 0.463 | 0.492 | 0.179 | 0.451 | 0.413 | 0.421 |
|  | GRNBoost2 | 0.537 | 0.397 | 0.474 | 0.515 | 0.267 | 0.501 | 0.491 | 0.455 |
|  | GENIE3 | 0.545 | 0.381 | 0.437 | 0.486 | 0.542 | 0.529 | 0.51 | 0.490 |
|  | Random | 0.505 | 0.498 | 0.507 | 0.512 | 0.62 | 0.468 | 0.51 | 0.517 |
| AUPRC | CellProphet | 0.605 | 0.572 | 0.761 | 0.611 | 0.282 | 0.592 | 0.308 | 0.533 |
|  | GRANGER | 0.066 | 0.073 | 0.083 | 0.154 | 0.025 | 0.026 | 0.017 | 0.063 |
|  | GRNBoost2 | 0.127 | 0.034 | 0.083 | 0.181 | 0.013 | 0.07 | 0.049 | 0.08 |
|  | GENIE3 | 0.137 | 0.042 | 0.078 | 0.183 | 0.028 | 0.061 | 0.063 | 0.085 |
|  | Random | 0.082 | 0.087 | 0.086 | 0.129 | 0.098 | 0.019 | 0.025 | 0.075 |
| F1 score | CellProphet | 0.43 | 0.689 | 0.832 | 0.677 | 0.337 | 0.586 | 0.394 | 0.564 |
|  | GRANGER | 0.134 | 0.133 | 0.152 | 0.247 | 0.054 | 0.038 | 0.034 | 0.113 |
|  | GRNBoost2 | 0.203 | 0.087 | 0.146 | 0.297 | 0.037 | 0.102 | 0.098 | 0.139 |
|  | GENIE3 | 0.218 | 0.108 | 0.162 | 0.322 | 0.042 | 0.108 | 0.112 | 0.153 |
|  | Random | 0.149 | 0.159 | 0.153 | 0.211 | 0.044 | 0.038 | 0.044 | 0.114 |
| EPR | CellProphet | 0.715 | 0.878 | 1.243 | 1.024 | 1.359 | 1.101 | 1.331 | 1.093 |
|  | GRANGER | 0.022 | 0.009 | 0.012 | 0.027 | 0.019 | 0.005 | 0.011 | 0.015 |
|  | GRNBoost2 | 0.311 | 0.075 | 0.14 | 0.354 | 0.142 | 0.158 | 0.291 | 0.21 |
|  | GENIE3 | 0.337 | 0.093 | 0.157 | 0.39 | 0.16 | 0.167 | 0.334 | 0.234 |
|  | Random | 0.221 | 0.142 | 0.148 | 0.24 | 0.17 | 0.057 | 0.128 | 0.158 |

## Notes

https://github.com/prsigma/CellProphet

### Competing Interest Statement

The authors have declared no competing interest.

### Summary of Updates

We have added results without using priors, included two additional competing methods on three datasets, and provided all corresponding results in tables. Furthermore, we have revised the title, abstract, introduction, and discussion sections.

## References

1. Bunne, C. et al. How to build the virtual cell with artificial intelligence: Priorities and opportunities. Cell 187, 7045–7063 (2024).

2. Badia-i-Mompel, P. et al. Gene regulatory network inference in the era of single-cell multi-omics. Nat Rev Genet 24, 739–754 (2023).

3. Bravo González-Blas, C. et al. SCENIC+: single-cell multiomic inference of enhancers and gene regulatory networks. Nat Methods 20, 1355–1367 (2023).

4. Aibar, S. et al. SCENIC: single-cell regulatory network inference and clustering. Nat Methods 14, 1083–1086 (2017).

5. Langfelder, P. & Horvath, S. WGCNA: an R package for weighted correlation network analysis. BMC Bioinformatics 9, 559 (2008).

6. Chen, L., Dautle, M., Gao, R., Zhang, S. & Chen, Y. Inferring gene regulatory networks from time-series scRNA-seq data via GRANGER causal recurrent autoencoders. Briefings in Bioinformatics 26, bbaf089 (2025).

7. Skok Gibbs, C. et al. High-performance single-cell gene regulatory network inference at scale: the Inferelator 3.0. Bioinformatics 38, 2519–2528 (2022).

8. Moerman, T. et al. GRNBoost2 and Arboreto: efficient and scalable inference of gene regulatory networks. Bioinformatics 35, 2159–2161 (2019).

9. Huynh-Thu, V. A., Irrthum, A., Wehenkel, L. & Geurts, P. Inferring Regulatory Networks from Expression Data Using Tree-Based Methods. PLoS ONE 5, e12776 (2010).

10. Wang, P. et al. Deciphering driver regulators of cell fate decisions from single-cell transcriptomics data with CEFCON. Nat Commun 14, 8459 (2023).

11. Kamimoto, K. et al. Dissecting cell identity via network inference and in silico gene perturbation. Nature 614, 742–751 (2023).

12. Wang, Y. et al. Reprogramming of regulatory network using expression uncovers sex-specific gene regulation in Drosophila. Nat Commun 9, 4061 (2018).

13. Vaswani, A. et al. Attention is All you Need. in Advances in Neural Information Processing Systems vol. 30 (Curran Associates, Inc., 2017).

14. Street, K. et al. Slingshot: cell lineage and pseudotime inference for single-cell transcriptomics. BMC Genomics 19, 477 (2018).

15. Tang, F. et al. mRNA-Seq whole-transcriptome analysis of a single cell. Nat Methods 6, 377–382 (2009).

16. Shojaie, A. & Fox, E. B. Granger Causality: A Review and Recent Advances. Annual Review of Statistics and Its Application 9, 289–319 (2022).

17. Browaeys, R., Saelens, W. & Saeys, Y. NicheNet: modeling intercellular communication by linking ligands to target genes. Nat Methods 17, 159–162 (2020).

18. Pratapa, A., Jalihal, A. P., Law, J. N., Bharadwaj, A. & Murali, T. M. Benchmarking algorithms for gene regulatory network inference from single-cell transcriptomic data. Nat Methods 17, 147–154 (2020).

19. The relationship between Precision-Recall and ROC curves. Proceedings of the 23rd international conference on Machine learning.

20. Aibar, S. et al. SCENIC: single-cell regulatory network inference and clustering. Nat Methods 14, 1083–1086 (2017).

21. Marucci, L. Nanog Dynamics in Mouse Embryonic Stem Cells: Results from Systems Biology Approaches. Stem Cells International 2017, 7160419 (2017).

22. Wang, Z., Oron, E., Nelson, B., Razis, S. & Ivanova, N. Distinct Lineage Specification Roles for NANOG, OCT4, and SOX2 in Human Embryonic Stem Cells. Cell Stem Cell 10, 440–454 (2012).

23. Raina, K., Dey, C., Thool, M., Sudhagar, S. & Thummer, R. P. An Insight into the Role of UTF1 in Development, Stem Cells, and Cancer. Stem Cell Rev and Rep 17, 1280–1293 (2021).

24. Shan, Y. et al. PRC2 specifies ectoderm lineages and maintains pluripotency in primed but not naïve ESCs. Nat Commun 8, 672 (2017).

25. Lavarone, E., Barbieri, C. M. & Pasini, D. Dissecting the role of H3K27 acetylation and methylation in PRC2 mediated control of cellular identity. Nat Commun 10, 1679 (2019).

26. Holtzinger, A., Rosenfeld, G. E. & Evans, T. Gata4 directs development of cardiac-inducing endoderm from ES cells. Developmental Biology 337, 63–73 (2010).

27. Oishi, Y. et al. SREBP1 Contributes to Resolution of Pro-inflammatory TLR4 Signaling by Reprogramming Fatty Acid Metabolism. Cell Metabolism 25, 412–427 (2017).

28. Varlakhanova, N. V. et al. myc maintains embryonic stem cell pluripotency and self-renewal. Differentiation 80, 9–19 (2010).

29. Liu, S. et al. NRF1 association with AUTS2-Polycomb mediates specific gene activation in the brain. Molecular Cell 81, 4663-4676.e8 (2021).

30. Zhu, Y. et al. Relaxed 3D genome conformation facilitates the pluripotent to totipotent-like state transition in embryonic stem cells.

31. Lieberman-Aiden, E. et al. Comprehensive Mapping of Long-Range Interactions Reveals Folding Principles of the Human Genome. Science (2009) doi:10.1126/science.1181369.

32. Buenrostro, J. D. et al. Single-cell chromatin accessibility reveals principles of regulatory variation. Nature 523, 486–490 (2015).

33. Zou, Z., Ohta, T. & Oki, S. ChIP-Atlas 3.0: a data-mining suite to explore chromosome architecture together with large-scale regulome data.

34. Robinson, J. T. et al. Integrative genomics viewer. Nat Biotechnol 29, 24–26 (2011).

35. Ohneda, K. & Yamamoto, M. Roles of Hematopoietic Transcription Factors GATA-1 and GATA-2 in the Development of Red Blood Cell Lineage.

36. Fujiwara, Y., Browne, C. P., Cunniff, K., Goff, S. C. & Orkin, S. H. Arrested development of embryonic red cell precursors in mouse embryos lacking transcription factor GATA-1. Proceedings of the National Academy of Sciences 93, 12355–12358 (1996).

37. Gutiérrez, L., Caballero, N., Fernández-Calleja, L., Karkoulia, E. & Strouboulis, J. Regulation of GATA1 levels in erythropoiesis. doi:10.1002/iub.2192.

38. Zhang, P. et al. PU.1 inhibits GATA-1 function and erythroid differentiation by blocking GATA-1 DNA binding. Blood 96, 2641–2648 (2000).

39. Howe, K. L. et al. Ensembl 2021.

40. Hayashi, T. et al. Single-cell full-length total RNA sequencing uncovers dynamics of recursive splicing and enhancer RNAs. Nat Commun 9, 619 (2018).

41. Nestorowa, S. et al. A single-cell resolution map of mouse hematopoietic stem and progenitor cell differentiation. Blood 128, e20–e31 (2016).

42. Shalek, A. K. et al. Single-cell RNA-seq reveals dynamic paracrine control of cellular variation. Nature 510, 363–369 (2014).

43. Chu, L.-F. et al. Single-cell RNA-seq reveals novel regulators of human embryonic stem cell differentiation to definitive endoderm. Genome Biol 17, 173 (2016).

44. Camp, J. G. et al. Multilineage communication regulates human liver bud development from pluripotency. Nature 546, 533–538 (2017).

45. Servant, N. et al. HiC-Pro: an optimized and flexible pipeline for Hi-C data processing. Genome Biol 16, 259 (2015).

46. Langmead, B. & Salzberg, S. L. Fast gapped-read alignment with Bowtie 2. Nat Methods 9, 357–359 (2012).

47. Reznikoff, W. S. Tn5 as a model for understanding DNA transposition.

48. Davies, D. R., Goryshin, I. Y., Reznikoff, W. S. & Rayment, I. Three-Dimensional Structure of the Tn5 Synaptic Complex Transposition Intermediate. Science (2000) doi:10.1126/science.289.5476.77.

49. Zhang, Y. et al. Model-based Analysis of ChIP-Seq (MACS). Genome Biol 9, R137 (2008).

50. Li, Z. et al. RGT: a toolbox for the integrative analysis of high throughput regulatory genomics data. BMC Bioinformatics 24, 79 (2023).

51. Rauluseviciute, I. et al. JASPAR 2024: 20th anniversary of the open-access database of transcription factor binding profiles.

52. Coifman, R. R. et al. Geometric diffusions as a tool for harmonic analysis and structure definition of data: Multiscale methods. Proceedings of the National Academy of Sciences 102, 7432–7437 (2005).

53. Coifman, R. R. & Lafon, S. Diffusion maps. Applied and Computational Harmonic Analysis 21, 5–30 (2006).

